# HID1 domain-containing protein 1 is required for normal cell proliferation in *Schizosaccharomyces pombe*

**DOI:** 10.64898/2025.12.02.691498

**Authors:** Anne-Philine Fritsche, Abdulrahman Alasmari, Mohammed Alshehri, Stephane Claverol, Heidi Fuller, Ramsay J. McFarlane, Mark A. Hooks

**Affiliations:** University of Bordeaux/CNRS, UMR 5200 Laboratory of Membrane Biogenesis, 71 Avenue Edouard Bourlaux, 33882 Villenave d’ Ornon, France; Bordeaux Imaging Centre, 71 Avenue Edouard Bourlaux, 33882 Villenave d’Ornon, France; University of Bordeaux Proteome, Functional Genomics Centre Bordeaux, Université de Bordeaux, 146 rue Leo Saignat 33076 Bordeaux Cedex, France; Faculty of Medicine and Health Sciences, Keele University, Keele, Staffordshire, UK ST5 5BG; North West Cancer Research Institute, Bangor University, Bangor, Gwynedd, LL57 2DG

**Keywords:** Cancer, Cancer Cell Line Encyclopedia, HID1 domain-containing protein, *Schizosaccharomyces pombe*, The Cancer Genome Atlas

## Abstract

HID1 domain-containing protein 1 (HID1) is a Golgi apparatus (GA) protein involved in the trafficking of cellular material. From reduced expression in cancer cell lines, it had been proposed to be a tumor suppressor in humans. In contrast, it is required for normal larval development of *Caenorhabditis elegans*, thereby raising an apparent contradiction for HID1 function regarding the inhibition or maintenance of cell proliferation. An extensive comparison of publicly available gene expression data revealed that *HID1* transcript levels in cancer cell lines were not representative of those in primary tumors, and the gene is amplified in many primary tumors. These findings question the role for its expression to suppress cell proliferation. Subsequently, we employed the model *S. pombe* to explore the role of HID1 in cell proliferation. The knock-out mutant *hid1Δ* exhibited a reduced proliferation phenotype due to a prolonged lag-phase but, eventually, attained parent strain rates of apparent proliferation. RNAseq-based transcriptomics revealed that *hid1Δ* retained features of metabolically stressed cells with quiescence-like transcriptional regulation. A label-free proteomic analysis revealed a lack of various membrane proteins suggesting cells with compromised Golgi apparatus (GA) function. The inability of *hid1Δ* to grow on agar containing a standard minimal medium suggested a link to metabolic deficiency. These findings point to a *hid1Δ*-related perturbation in GA function that compromises cell recovery from metabolic change.

## Introduction

The Golgi apparatus (GA) is the primary organelle for the sorting and trafficking of lipids and proteins within a cell and to the cell exterior. The requirements of the cell to coordinate exocytic and endocytic processes with intracellular trafficking of proteins and lipids is why the GA is a hub linking such mechanical actions with signalling processes (Mayinger, 2011; Makowski 2017). Given the wide variety of functions of the GA, it is not surprising that disrupting this organelle in humans has wide-ranging consequences, such as severe morphological abnormalities (El Ghouzzi et al. 2003, Dupuis et al. 2003), cognitive disease (Ayala and Colanzi, 2017), and tumorigenesis (Sechi et al. 2020). In humans, HID1 is a protein of the *trans*-Golgi network (Hummer et al. 2023). It has been observed also in the medial-GA, Dense Core Vesicles and in the plasma membrane of neuron synapses (Wang et al. 2011; Hummer et al. 2017). Protein localization studies have implied a role in the trafficking of material from the inner membrane system to the cell surface and cell exterior (Du et al. 2016; Mesa et al. 2011; Yu et al. 2011), and the loss of HID1 in rat cells affects protein excretion and intracellular trafficking (Hummer et al. 2017). The loss of *Caenorhabditis elegans* HID1 in certain types of neurons can alter responses to extracellular cues and thus perturb normal physiological functions (Sharabi et al. 2014) and mutations in HID1 have been linked to severe genetic diseases in humans (Schänzer et al. 2021).

HID1 was reported initially as a potential tumor suppressor referred to as DOWN-REGULATED in MULTIPLE CANCERS 1, whereby the expression of *HID1* was reduced or lacking in a substantial proportion of cancer cells lines used in the study (Harada et al. 2001). Subsequently, *HID1* was found to comprise part of a core of discriminatory genes for pituitary cancer (Aydin and Arga, 2019). However, the gene encoding the HID1 ortholog of *C. elegans* was isolated based on a screen to identify genes regulating the switch of larvae into Dauer development upon change in temperature (Ailion and Thomas, 2003). The observation that Dauer formation was initiated under a temperature conducive for normal reproductive development suggested that reduced expression could reduce apparent cell proliferation and induce quiescence. Great strides have been made to elucidate the structure and function relationships of HID1 in animals (Hummer et al. 2023), but the relationship between *HID1* expression and its role in cell proliferation remains to be resolved.

During database searches for genes potentially involved in tumorigenesis (Feichtinger et al. 2012; McFarlane and Wakeman, 2017), we observed *HID1* to exhibit highly variable, tissue-type dependent gene expression, including up-regulation in some primary tumors. This raises the question if cancer cell lines represent an appropriate model for gene expression and functional characterization for *HID1* in primary tumors. We present a comprehensive comparison of transcript levels in cancer cell lines compared to primary tumors to address this question and explain the conflicting findings in between the human and worm work. Subsequently, we employed the genetically tractable model organism, *Schizosaccharomyces pombe*, to investigate the role of HID1 in cell proliferation. The genus *Schizosaccharomyces* is unique in that it contains three distinct orthologs of human HID1 (Wood et al. 2002; Hooks et al. 2025), of which the paralog Hid1 of *S. pombe* has been localized to the GA (Kouranti et al. 2010). Previous work has shown that the deubiquitinase Upb5 is mis-targeted in the mutant lacking Hid1 (*hid1Δ*), due to either the loss of direct interaction of Ubp5 with Hid1 (Kouranti et al. 2010) or altered GA structure (Hooks et al. 2025). We find that *hid1Δ* has a prolonged lag-phase, but can attain normal rates of proliferation. Even at maximal proliferation rate, it retains transcriptional characteristics of cellular quiescence that are likely caused by perturbed protein trafficking and metabolism.

## Materials and Methods

### Computation and statistical analyses

The list of programs, hardware platforms and databases used in this study is given in Supplementary Table 1.

### Comparison of transcript amounts among cancer datasets

The transcript counts by gene for cancer cell lines, normal tissue and primary tumors for the various tissue types were obtained from repositories of the Cancer Cell Line Encyclopedia (CCLE,) and The Cancer Genome Atlas (TCGA) on February 12, 2019. The CCLE dataset and TCGA datasets had been annotated under human genomes GRCh37 and GRCh38, yielding counts for 56202 and 60483 unique transcript entities, respectively. The CCLE data was combined with the normal and tumor data based on the tissue of origin for subsequent normalization using DESeq2. Normalization was conducted only using genes common to both cell lines and tissues. In brief, since the versions of human genomes and Ensembl annotations differed between cell lines and tissues, the Ensembl gene identifiers for the cell line datasets were stripped of the gene version number and the core gene identifiers used to retrieve the corresponding genes and their counts from the TCGA datasets. The resulting number of gene entities for normalization for each tissue type was 53196. The final number of genes in each set was slightly different after removal of genes for which there were no counts present in any of samples. The 22 Recurrent tumor samples among the TCGA data sets were removed and, although added to appropriate plots, a statistical analysis was not conducted on metastatic samples due to low sample numbers. In total, there were 19 RNAseq-based transcriptomic projects to which we could attribute normal tissue, primary tumor and cell line data. Celligner quantitative and meta data were downloaded from FigShare (Data ref: https://figshare.com/articles/dataset/Celligner_data/11965269; Warren et al. 2021).

### Generation and molecular characterization of *hidΔ* mutants

Mutant strains were generated from a common multi-auxotrophic strain (BP90: *h*^-^ *ade6-M26 leu1-32 ura4-D18*) by recombination-mediated gene replacement (Bähler et al. 1998). PCR products used for transfection were generated using the template pFA6a-NatMX6 and selection for positive transformants was conducted using the cloneNat (Werner Bioagents, Jena, Germany). The list of strains employed in the study is presented in Supplementary Table 2. DNA was extracted using the Wizard Genomic DNA Extraction kit (Promega Corp., United Kingdom) according to the manufacturer’s instructions for yeast DNA isolation. Total RNA for RT-PCR was extracted by the hot phenol/chloroform method according to (Lyne et al. 2003). First-strand cDNA was produced by random-priming from 1 μg of total RNA using the QuantiTect^TM^ Reverse Transcription kit according to manufacturer’s instructions (QIAGEN Ltd., United Kingdom). PCR was conducted on a MJ Research thermocycler with the following cycling parameters: an initial denaturation step of one cycle of 1 min at 94°C, 34 cycles of 15 sec at 94°C, 15 sec at 55°C, and 15 sec at 72°C, followed by an extension period of 72°C for 5 min. The list of marker and gene/genome-specific primers used for genotyping and RT-PCR is given in Supplementary Table 3. For the complementation analysis, Cntl and *hid1Δ* strains were transformed with the empty vector pREP41_ccdB2 (Matsuyama et al. 2008) and *hid1Δ* with the same vector containing the *hid1+* gene. Transformation was conducted using a standard LiAc-based protocol and selection was done on solid agar medium without leucine.

### Cell culture and sample preparation

All strains were incubated in 5 ml YE media (0.5% w/v yeast extract, 3% w/v glucose), containing the standard auxotrophic supplements where required (YEL), at 30°C with shaking 200 RPM to a cell density of 5×10^7^ cells·ml^-1^. Cells were diluted in 5 ml of fresh YEL to a cell density of 5×10^5^ cells·ml^-1^. Cultures were incubated as above for 12 h before aliquots were taken as desired times up to 24 h. Generation times under stress treatments were calculated from cell counts taken at 12h after dilution. Total cell counts were obtained using the Nexcelom Bioscience Cellometer Mini (Ozyme, Yveline France). Real-time monitoring of culture turbidity was conducted using an Oy Growth Curves Ab Ltd. Bioscreen C**™** (Thermo Fisher Scientific, France) operating at 30° C with continual shaking at maximum speed. Absorbance readings at 600 nm were recorded every 10 min following a 5 s pause in shaking. Typical well volumes for honeycomb plates were 150 μl with an initial O.D. at 600 nm of 0.05. For the complementation analysis, all strains were grown over-night in YE without leucine and diluted into the same medium at a cell density of 5×10^5^ cells·ml^-1-^. Cultures were incubated for 16h at 30°C with shaking at 200 RPM. Aliquots of cell culture were counted for cell amounts using a standard haemocytometer. For post-genomic analyses, control and mutant strains were grown in 50 ml YEL media with shaking at 200 RPM at 30°C to a cell density of 5×10^7^ cells·ml^-1^. Cells were harvested by centrifugation, washed rapidly in 10 ml H_2_0, re-centrifuged and the pellets frozen in liquid N_2_. Pellets from two independent strains of each mutant genotype were combined and powdered while frozen using a Freezer/Mill 6750 (SPEX Europe, United Kingdom). These combined samples were divided and used for both transcriptomic and proteomic analyses.

### Transcriptomic analyses

Total RNA was extracted from approximately 50 mg of powdered cells for each of the 11 samples according to Lyne et al., 2003. In brief, the extraction protocol involves an initial purification of RNA by phenol/chloroform extraction followed by further purification using the RNAeasy^TM^ RNA extraction kit (Qiagen). The quality of the RNA was determined on an Agilent 2100 bioanalyzer (Agilent, France) using a RIN threshold of 0.9. The RNA was depleted of ribosomal RNAs prior to creation of the RNAseq libraries. RNAseq was performed on an Illumina MySEQ^TM^ to obtain a total of 21.1 million reads total for the 11 samples. The complete genome and corresponding transcriptome files were downloaded from Pombase (www.pombase.org, modified 2018-09-04). Mapping and read quantification of the paired-end fastq files were done with Salmon v 1.0.0. against the *S. pombe* transcriptome and employing the full genome as decoy (Supplementary Table 4), and against the full genome using STAR (Supplementary Table 5). Normalization of gene counts across samples and determination of differential gene expression based on consensus counts were done using DESeq2. The only pre-filtering was done prior to normalization and consisted of the removal of those genes with zero counts across all samples, The internal filtering employed for differential expression by DESeq2 appears strict for limiting false positives (Soneson and Robinson 2018).

### Label-free protein quantification

Approximately 100 mg of frozen powder was further homogenized using a Qiagen Tissuelyser II in a 2 ml screw-cap tube containing acid-washed glass beads (Sigma-Aldritch, France) and 0.4 ml of homogenization buffer (25 mM MOPS (pH 7.2), 60 mM β-glycerophosphate, 15 mM p-nitrophenylphosphate, 15 mM MgCl_2_, 15 mM EGTA, 1 mM DTT, 0.1 mM sodium vanadate, 1% Triton X-100, 1 mM PMSF, 1 x Yeast Protease Inhibitor Cocktail plus 1 x Mammalian Protease Inhibitor Cocktail from Sigma-Aldritch, France). After clarification by centrifugation, protein content was measured using the Pierce**®** BCA^TM^ method according to published instructions (ThermoFisher Scientific). Visualization of total protein extracts was done by discontinuous SDS-PAGE using 5% and 10% w/v acrylamide for stacking and separating gels, respectively. Protein preparation for label-free proteomics entailed loading 10 μg of total protein from each sample onto a 10% SDS-PAGE and allowed to migrate until all protein had just entered the gel. Proteins were visualized by staining with colloidal blue (Biorad, France) and from each lane a gel slice containing the protein was excised. Proteins were digested in gel using trypsin and the peptides eluted from the gel. The samples were acidified, injected into a Q-Exactive^TM^ quadripole Orbitrap mass spectrometer (ThermoFisher Scientific) and the data collected over a liquid chromatography run period of 120 min. The resulting mass spectra were interrogated with Proteome Discoverer^TM^ v 1.4 (ThermoFischer Scientific, France) using the *S. pombe* proteome as template for identifying peptide masses. Proteins changing significantly between mutant and control genotypes were determined by ANOVA (threshold, *p-value* < 0.05) and only proteins with a minimum of two peptides were quantified for differential amounts.

## Results

### Cell lines do not represent HID1 expression in primary tumors

The proposal that reduced expression of *HID1* leads to cell proliferation was based on an RT-qPCR analysis of a limited number of cancer cell types and restricted to immortal cell lines (Harada et al. 2001). From this report, a role in cancer development remains a prominent feature of its functional description in databases, such as UniProt (https://www.uniprot.org). Publicly available transcriptome data from both microarray and RNAseq experiments, comprising a wide variety of cancer types, allows direct investigation of the correspondence between cell line and tumor expression compared to that in disease-free normal tissues. Cell line transcript data for RNAseq-based experiments was retrieved from the CCLE and the corresponding normal and tumor data from TCGA. The gene counts for the cell lines tumors and corresponding normal tissues were grouped by cancer type and normalized using DESeq2. This has enabled us to conduct an analysis of the expression of individual genes in cancer cell lines compared to tumors. For every cancer type characterized by RNAseq, transcript levels for *HID1* in cell lines were less than for the corresponding tumors, including inverted median differences in transcript amounts for nine of the 19 projects investigated (Fig. 1a; Supplementary Fig. 1).

**Figure 1.**
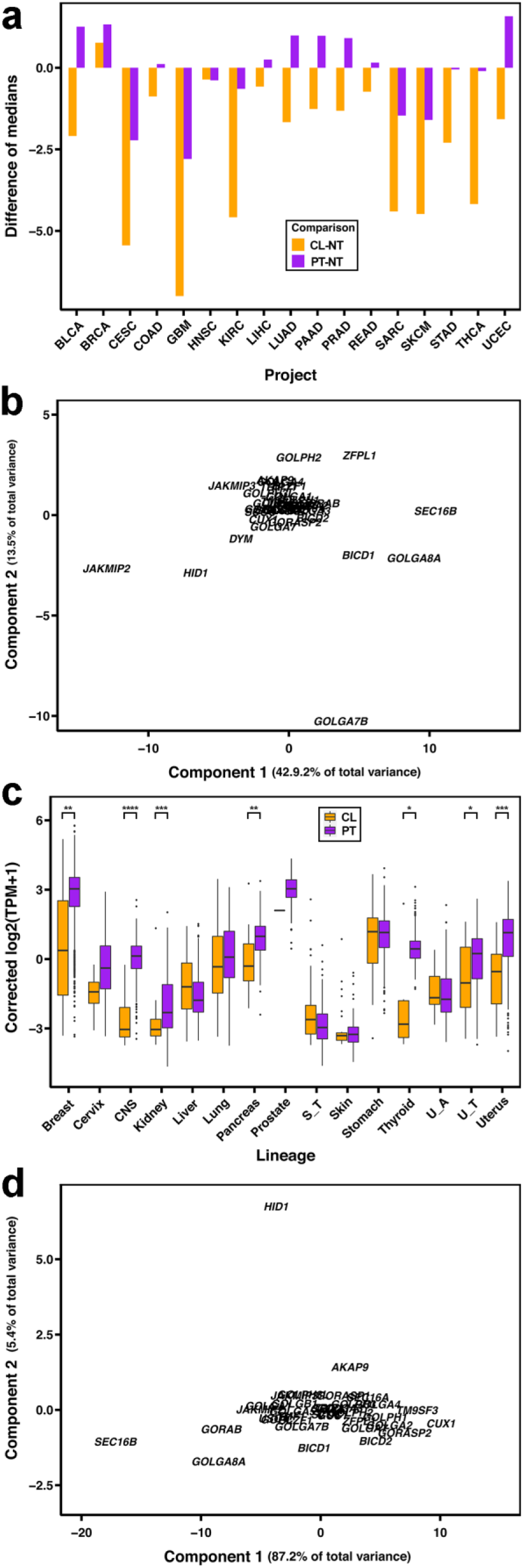
Comparison of *HID1* transcript amounts between primary tumors and cancer cell lines. The test for significance of medians in Figure S1 and Figure 1C was conducted using Mood’s Median Non-parametric Hypothesis Test. The number of samples for each tissue type per project is given in the information file for the normalized values on the Open Science Framework (see data availability below). **(A)** Values are presented as the difference in the log_2_-adjusted median values of *HID1* transcript counts for the data presented in Figure S1. **(B)** PCA scores plot of the median transcript amounts for Golgi Protein-encoding Genes as shown in Figure S2. The scores were calculated for each gene using the medians for both the cell lines and primary tumors across all projects. **(C)** Boxplot presentation of Celligner-corrected expression values for cell lines and primary tumors according to tissue of origin. The dots represent outliers from quartiles represented by the boxes and whiskers. The asterisks represent the FDR-adjusted P values: * < .05; ** < .01; *** < .001; **** < .0001. **(D)** PCA scores plot of median transcript amounts for Golgi Protein-encoding genes as shown in Figure S3. The scores were calculated for each gene using medians for both cell lines and primary tumors across all projects. Abbreviations: CL, cell line; NT, normal tissue; PT, primary tumor.

The trend of lower expression values in cell lines did not appear to be a general phenomenon. In order to determine if our normalization method yielded artificially lower transcript levels for cell lines, we investigated the expression of a cohort of 28 genes encoding GA proteins. This cohort was comprised of the genes described in Barlow et al., 2018, including orthologs, and other genes that have been implicated in cancer development, such as *GOLPH3L* and *GOPC*. Four of the genes, *AKA9*, *GOLGA5*, *GOPC*, and *TRIP11*, are mentioned in TCGA as potential cancer drivers. After normalization, most genes had median transcript levels in both cell lines and tumors across cancer types that were indistinguishable from those in normal tissue (Supplementary Fig. 2). Several genes, including *HID1*, exhibited either variable transcript levels across cancer types and/or a systematic difference between cell lines or tumors when compared to normal tissue. *HID1*, *DYM*, *JAKMIP2, JAKMIP3* trended toward fewer transcripts in cell lines, whereas *BICD1*, *GOLGA8A*, *SEC16B* and *ZFPL1* exhibited slightly greater transcript levels. The deviation of these genes from the cluster of adjustable genes was clearer when the median values were subjected to a principal component analysis (Fig. 1b). Genes with median cell line values less than tumor or normal tissues were projected negatively across PC1, whereas those genes with higher cell line values were positive. This normalization procedure does not appear to alter artificially the fundamental expression behavior of genes as shown by the median transcript levels of two supplementary sets of genes (Supplementary Fig. 2). One set was comprised of some well-characterized transcription effectors, which do not show altered transcript levels in tumors, and another set of genes induced ubiquitously in tumors (portal.gdc.cancer.gov). The median transcript levels of the transcriptional effectors varied little, if any, and those for the induced genes were greater in both cell lines and tumors, thereby demonstrating that our normalization process did not introduce a systemic bias into our adjusted transcript amounts.

The recently reported Celligner employs a PCA-based approach to normalize CCLE cell line and TCGA tumor transcript data and has proven useful to identify cell lines that do not match for a particular tumor type (Warren et al. 2021). Although Celligner was not designed to investigate systematic differences in transcript levels for individual genes among cell lines and tumors, as per the approach described here, it may be suitable to identify gene outliers that do not conform to global correction. The transcript amounts for the gene sets described above were taken from the Celligner data for calculation of median values. *HID1* transcript levels were less in cell lines than tumors for 10 of the 15 tissue types reported (Fig. 1c). Celligner abolished differences among cell line and tumor transcript amounts for virtually every other gene investigated (Supplementary Fig. 3). *HID1* was the notable exception and accounted for the variance represented by the second principal component of a PCA scores plot of cell line and tumor median transcript values (Fig. 1d). Therefore, the Celligner approach reinforced the conclusion from our initial assessment that *HID1* expression in cell lines is not indicative of expression in tumors.

In contrast to the cell lines, we found that *HID1* transcripts were greater in tumor samples than normal tissue samples for 11 of the 19 TCGA projects analysed (Fig. 1a). This could be explained by copy number amplification as suggested by tumor sequencing data from TCGA. The global mutation frequency of *HID1* is low compared to some well-known transcriptional effectors, such as *TP53*, *PTEN*, *FOXO1* and *FOXO3,* and *MTOR* (Fig. 2a). Although the number of mutations within *HID1* was greater than the median of the cohort of all genes, it fell well within the spread of mutation number observed for the cohort genes for GA proteins, including the potential cancer drivers *AKAP9* (Hu et al. 2016), *GOLGA5* (Longo et al. 2016), *GOPC* (Johnson et al. 2017), and *TRIP11* (Popławski et al. 2017). In addition, *HID1* was well below the median for copy number loss relative to the cohort of all genes, the transcriptional effectors, and the cohort of genes encoding GA proteins (Fig. 2b). In contrast, copy number gain for *HID1* was well above median for all genes, and only the potential cancer driver *GOLPH3L* (Feng et al. 2017) was revealed to have a higher incidence of copy gain than *HID1* for the set of genes examined (Fig. 2c). Although our data suggests that decreased *HID1* expression does not play a general role in abnormal cellular development leading to cancer, much remains to be elucidated regarding its role in metabolic maintenance and cell proliferation. This is particularly so for single-celled eukaryotes, where HID functions remain virtually unknown.

**Figure 2.**
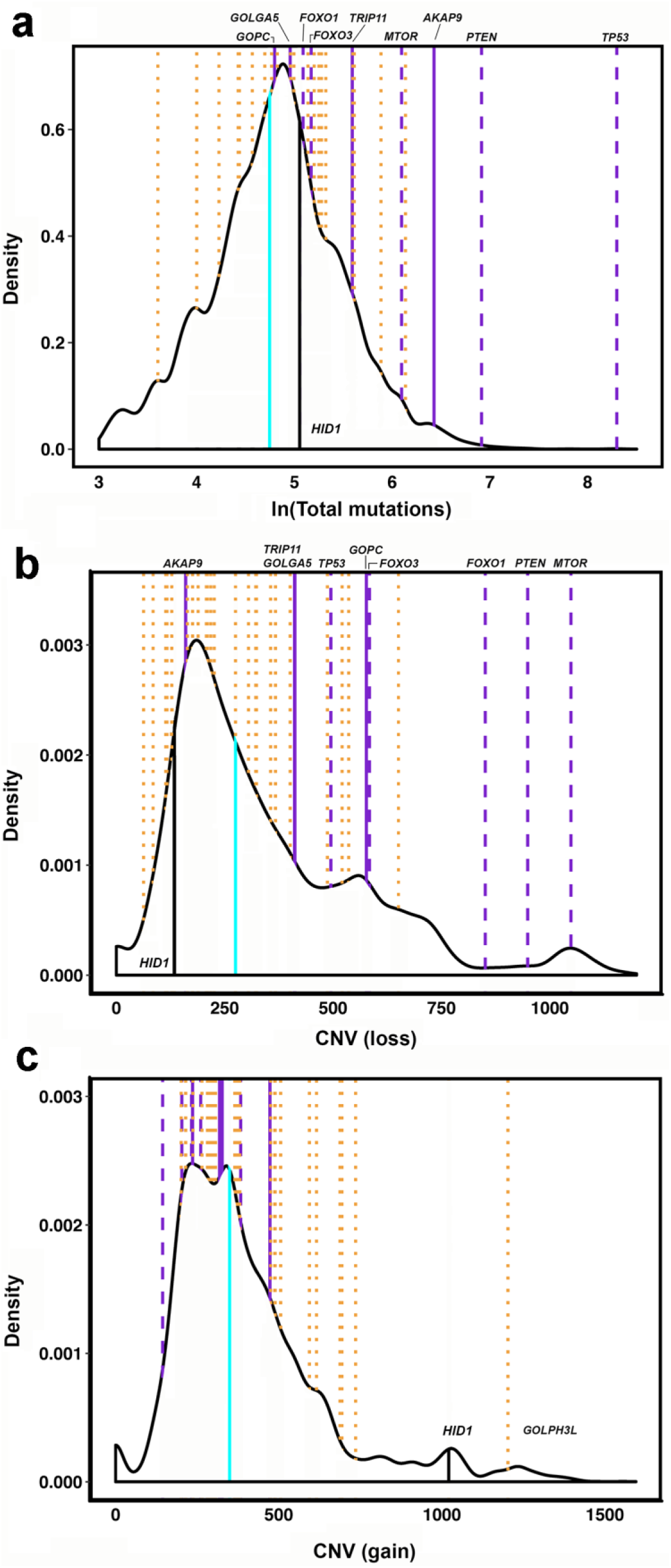
Density plots of copy number variation across TCGA samples. Mutation number and CNV values for all genes across the TCGA projects were downloaded from TCGA. The cohort was represented by 10198 cases. The total number of variations across the cohort was 10741. Vertical line designations: solid cyan, median number of entities.; solid black, HID1; dotted orange, Golgi protein-encoding genes; solid purple, CGC genes encoding Golgi proteins; dashed purple, examples of CGC non-Golgi protein encoding genes, lines. **(A)** Total number of mutations per gene in cohort. The number of total mutations has been log (natural) transformed for value separation. Only the Golgi and non-Golgi CGC genes have been labelled. **(B)** Number of samples in the cohort with the gene lost. Only the Golgi and non-Golgi CGC genes have been labelled. **(C)** The number of samples in the cohort for which the gene copy number was increased.

### Hid1 deficiency delays onset of apparent maximum proliferation rate

Deletion mutants were created for which each of the three *hid* paralogs for comparative physiological and molecular characterization. The mutants were tested for some traits already reported from large phenotypic screens to ensure the validity of subsequent experimental observations. However, there is little comparative information for *hid2Δ* than for the other mutants, because *hid2Δ* is not present in the KRIBB-BiONEER-CRUK consortium mutant collection, commonly used for large phenotypic screens (Kim et al. 2010). The mutant *hid1Δ* has been shown to be unaffected by Brefeldin A (Hooks et al. 2025), which confirmed previous reporting of Brefelding A resistance (Rodríguez-López et al. 2023), but highly sensitive to the ArfGEF inhibitor Golgicide A (Hooks et al 2025). The mutant *hid1Δ* has been reported to be sensitive to salt stress (Kennedy et al. 2008), which we confirmed, and observed that *hid2Δ* may be slightly tolerant (Supplementary Fig. 4a). Although *hid3Δ* had been reported as resistant to H_2_0_2_ at 0.5 mM, it was not reported as resistant at 1 mM (Rodríguez-López et al. 2023). We observed sensitivity of both *hid1Δ and hid3Δ*, greater even than the control *rad3Δ*, at a concentration of 0.7 mM (Supplementary Fig. 4b). Thiabendazole (TBZ) has been shown to disrupt the GA in *S. pombe* (Ayscough et al. 1993), therefore, we tested it on the proliferation of the *hid* KO mutants. All mutants were insensitive at a concentration that affected the control strain *bub1Δ*, a KO mutant of the mitotic spindle checkpoint kinase (Supplementary Fig. 4c; Bernard et al. 1998). Although, we obtained some apparent discrepancies with other reports, the KO mutants exhibited responses to agents that are consistent with potential disruptions to GA function and subsequent protein trafficking.

The increase in generation time that had been observed for *hid1Δ* in the absence of any stress agent, suggested a fundamental effect of the lack of Hid1 on proliferation rate (Hooks et al 2025). We investigated this in more detail by counting cell numbers during the culture of strains from dilution to stationary phase in standard YEL medium. In addition to the parental strain BP90, two independent isolates of strain MH7 (Cntl), for which the antibiotic marker, apparently, had inserted into another part of the genome, were included primarily to serve as controls for the subsequent quantification of profiling data. Fitting of the resulting curves to a logistic growth function allowed for quantification of the four critical apparent proliferation parameters: the initial cell concentration, the maximal rate of proliferation, the carrying capacity of the culture, and the lag phase leading to exponential proliferation. The mutant *hid1Δ* took longer to reach a particular cell density than BP90, Cntl and other mutant strains (Fig. 3). This effect was specific for the loss of Hid1, because the transcript levels of the other two paralogs within the *hid1Δ* mutant appeared unaffected (Supplementary Fig. 5). From the four parameters calculated, the primary difference between *hid1Δ* and the other strains was the lag phase, where it was estimated to have entered exponential growth approximately three hours later than BP90. The apparent rate of cell proliferation of *hid1Δ* at mid-log phase was equivalent to those of the other genotypes. The delay in attaining maximum proliferation rate was not due to a reduced viability of *hid1Δ* cells (Supplementary Fig. 6). The *hid1Δ* phenotype was specifically due to loss of Hid1 as shown by retrieval of Cntl-level cell counts by expression of *hid1+* in the mutant (Fig 3, inset). The question raised by this observation was if changes in transcript and protein were consistent with a reduced apparent proliferation of cells. We addressed this question by analysing the transcriptome and proteome of the various strains harvested at the calculated midpoints from the cell-count plots. Choice of this time point provided a direct comparison of the various strains at an equivalent stage of development.

**Figure 3.**
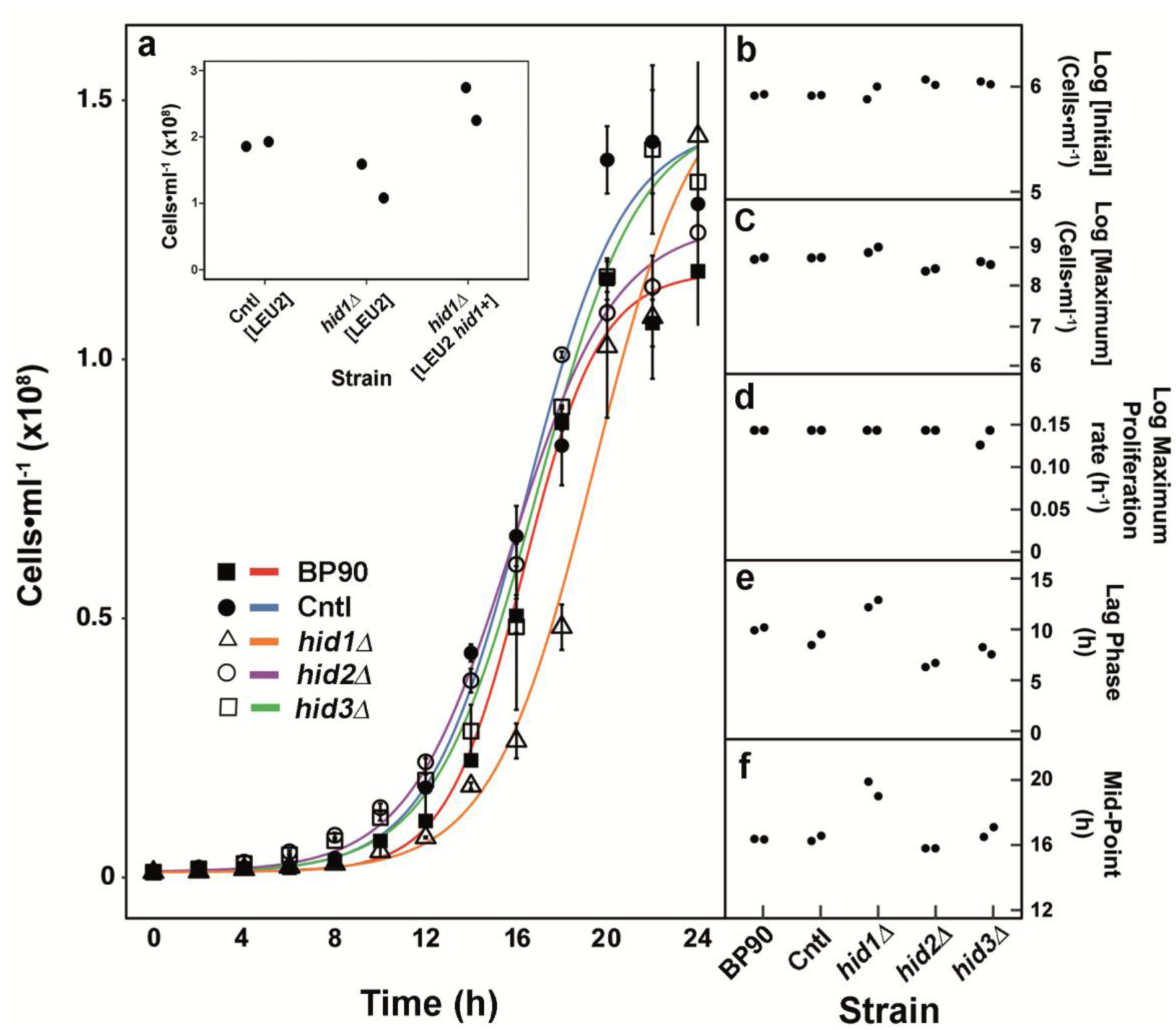
Apparent proliferation of *hid* mutants. All strains were propagated in liquid YE with supplements as indicated in Materials and Methods. **(A)** Culture turbidity profiles of two independent mutant strains based on colony counts from cultures taken every two h for 24 h. For clarity of presentation, the symbols and bars represent the mean and the range, respectively, of the cell count values from the two different isolates for each genotype. **(Inset to A)**. Cell counts of mutant *hid1Δ* expressing *hid1+* compared to CNTL and *hid1Δ* carrying the empty vector (*LEU2*-based selection). **(B-F)** The colony-count values were log-transformed to obtain the growth parameters shown in panels (**B-E**) as determined using the Baranyi-Roberts Growth Model from the R package ‘growthrates’. Curves were fitted to average values using the four-parameter logistic function from R package ‘drc’ to obtain the mid-point value for each curve **(F)**. For panels **(E-F)**, the ranges of points for *hid1Δ* do not overlap with those of the other strains.

### Hid1 deficiency causes quiescence-like transcriptional changes

Since only *hid1Δ* presented the slow proliferation phenotype, it was the primary target for molecular characterization. BP90 harboring the antibiotic resistance gene (strain MH7) was used for quantifying differential expression. BP90 and *hid3Δ* (MH11) were included for evaluating biological effect. Our analysis comprised three technical replicates of each strain, except for the two BP90 samples, with biological variation accounted for by mixing equal quantities of RNA from two corresponding isolates (Supplementary Fig. 7). For both STAR and SALMON, sample normalization and DE gene expression were performed using DESeq2. Investigation of the *hid+* genes themselves indicated that *hid1+* and *hid3+* transcript levels were similar, and both were greater than those for *hid2+*. Transcript amounts for *hid1+* and *hid3+* were above, or at least, at median levels compared to the total genes when mapped to the genome and transcriptome, respectively (Supplementary Fig. 8a, 8b). The relative differences in *hid+* expression in wild-type and Cntl strains were confirmed by RT-qPCR, although the amount of *hid2+* transcript was less than expected based on the RNAseq data (Supplementary Fig. 8c).

The quantity of reads obtained in the study was sufficient to allow quantification of transcript levels and comparison among samples, since both low and high number of input reads did not appear to bias the clustering of samples according to the principal component analysis (PCA) of normalized transcript counts (Supplementary Fig. 9). According to the PCA, the Cntl, BP90 and *hid3Δ* samples clustered together with only minor spread across PC2 (11% of the variance), suggesting little difference in transcript levels. In contrast, *hid1Δ* was removed from the others with two samples well displaced across PC1 and another displaced across PC2. Therefore, it was expected that *hid1Δ* would show the greatest number of differentially expressed genes. Using a standard *p*-adjusted value of 0.05, STAR produced 392 UR and 266 DR for *hid1Δ*. At the same *p*-adjusted value, SALMON yielded less than 40 % (153 UR, 99 DR) of the total DE genes obtained by STAR for *hid1Δ*. In order to compensate for the low DE gene output, the DESeq2 *p*-adjusted value for the SALMON mapping was increased to 0.1. Adjustment yielded 244 UR and 167 DR genes, of which 206 UR and 126 DR were present in the STAR output of DE genes (Supplementary Fig. 10a, 10b). Both STAR and SALMON yielded a very low number of DE genes for *hid3Δ* and BP90, with the most being 29 DR for *hid3Δ.* There was also substantial overlap between *hid3Δ* and BP90 DE genes (Supplementary Fig. 10a, 10b). The number of DE genes yielded by each of the mutants showed that the lack of Hid1 exhibited substantial transcript level changes compared to the lack of Hid3. The low number of DE genes for *hid3Δ* (Supplementary Table 6), upon removal of those common to BP90 (Supplementary Table 7), did not provide a cohort large enough for any statistically significant over-represented gene ontological (GO) classes.

In order to preserve a maximum of information of transcriptional changes for the GO analysis of *hid1Δ*, we took the union of the STAR and SALMON DE genes, which gave a total of 428 UR and 291 DR genes (Supplementary Fig. 10b). Interestingly, the loss of Hid1 resulted in a net increase in DE genes, but the degree of changes to transcript amounts was not dramatic with the greatest change being < 10-fold (Table 1). Evident within the cohort of UR genes were those encoding plasma membrane transporters and other cell-surface proteins, whereas the most highly DR genes did not appear to belong to any particular class (Table 1; Supplementary Table 8). UR and DR genes were subjected to gene enrichment analysis (EA) using PANTHER and AnGeLi to identify those processes potentially affected by the changes in gene expression. The analysis was conducted on the protein-coding genes using the full transcriptome as reference and both programs gave similar results. In addition to commonly observed effects on ribosomal processes and protein synthesis, the UR genes highlighted the altered expression of genes encoding integral membrane proteins and the regulation of gene expression (Supplementary Table 9). The latter primarily reflects the change in transcript amounts of genes related to transcription by RNA polymerase II. Also significantly enriched were genes encoding proteins related to nuclear transport of proteins and enriched cellular component genes related to ribosome biogenesis. Under-represented GO groups related to membrane-associated proteins and mitochondrial proteins. The DR genes were characterized primarily by a decrease in mitochondrial energy producing capacity with decrease gene expression for oxidative phosphorylation, small molecule catabolism, electron transfer and mitochondrial metabolic processes (Supplementary Table 10). GO classes also DR were related to organellar membrane proteins. EA using AnGeLi provided a means to compare the effect of expression changes of our mutants to those of quiescent cells from the study of Marguerat et al., 2012. For the *hid1Δ* UR genes, 163 and 202 genes were greater than the mean mRNA copy number for proliferating and quiescence gene sets, respectively, with 142 genes common to both proliferating and quiescent gene sets. EA revealed that common genes were dominated by those highly expressed glycolysis and cytoplasmic translation. EA did not yield significant GO classes for the 21 genes unique to the proliferating set. However, EA of the 60 genes unique to the quiescent dataset revealed enrichment for the GO classes relating to RNA polymerase II-related regulation of gene expression, response to starvation, and cellular response to nutrient levels (Supplementary Table 11). EA relating to molecular function highlighted RNA polymerase II transcription factor activity. The *hid1Δ* DR genes were under-represented in relation to mRNA copy number for both proliferating and quiescent cells, and there was extensive overlap between the two sets of the low mRNA copy number.

**Table 1.** Top DE genes in *hid1Δ*. The genes shown have been calculated by DESeq2 as significantly UR or DR by at least 2-fold. There were another 63 gene DR greater than 2-fold.

| Name | Log<sub>2</sub>_fold | Description |
| --- | --- | --- |
| **UR** |  |  |
| *gsf2* | 2.98 | cell surface glycoprotein, galactose-specific flocculin |
| *fip1* | 2.86 | plasma membrane iron transmembrane transporter |
| *SPAC27D7.09c* | 2.12 | But2 family protein, similar to cell surface molecules |
| *pfl3* | 1.83 | cell surface glycoprotein, flocculin |
| *SPBPB10D8.01* | 1.80 | cysteine transmembrane transporter |
| *gst2* | 1.64 | glutathione S-transferase |
| *caf5* | 1.62 | plasma membrane spermine family transmembrane transporter |
| *SPBC9B6.03* | 1.24 | zf-FYVE type zinc finger protein, involved in endosomal transport |
| *cys2* | 1.23 | homoserine O-acetyltransferase |
| *SPBC409.08* | 1.21 | spermine family transmembrane transporter |
| *met1* | 1.17 | uroporphyrin methyltransferase Met1 |
| *bfr1* | 1.14 | plasma membrane brefeldin A efflux transporter |
| *sfp1* | 1.14 | DNA-binding transcription factor |
| *cbp1* | 1.10 | CENP-B homolog |
| *mpe1* | 1.10 | mRNA cleavage ubiquitin-protein ligase E3 |
| *SPAC2H10.01* | 1.10 | DNA-binding transcription factor zf-fungal, binuclear cluster type |
| *adh8* | 1.05 | alcohol dehydrogenase |
| *dfr1* | 1.05 | dihydrofolate reductase/lysophospholipase fusion protein |
| **DR** |  |  |
| *urh1* | 4.81 | uridine ribohydrolase Urh1 |
| *SPAC977.18* | 2.98 | conserved fungal plasma membrane protein |
| *SPBPB2B2.18* | 2.96 | conserved fungal plasma membrane protein |
| *ddr48* | 2.86 | DNA damage-responsive protein |
| *SPCC70.08c* | 2.75 | methyltransferase |
| *uck2* | 2.24 | uracil phosphoribosyltransferase |
| *smp4* | 2.21 | Stm1/Oga1 family protein Smp4 |
| *urg2* | 1.98 | uracil phosphoribosyltransferase |
| *urg1* | 1.94 | GTP cyclohydrolase II |
| *ksh1* | 1.70 | Golgi kish family protein Ksh1 |
| *SPAC521.03* | 1.67 | short chain dehydrogenase, human DHRS7 family |
| *vps66* | 1.62 | 1-acylglycerol-3-phosphate O-acyltransferase |
| *put1* | 1.56 | proline dehydrogenase |
| *SPAC29B12.13* | 1.49 | CENP-V, S-(hydroxymethyl)glutathione synthase |
| *emi5* | 1.48 | succinate dehydrogenase complex assembly protein |
| *nud3* | 1.46 | nuclear distribution protein NUDC homolog |
| *hem13* | 1.41 | coproporphyrinogen III oxidase |
| *dap1* | 1.39 | cytochrome P450 regulator |
| *aim41* | 1.38 | mitochondrial aspartyl/glutamyl-tRNA amidotransferase subunit B-related |
| *urg3* | 1.38 | DUF1688 family fungal conserved protein, uracil or riboflavin metabolism |

The relationship between gene expression and cellular quiescence for the *hidΔ* mutants was investigated further by employing our quantification pipeline to analyse the cell proliferation and quiescence RNAseq data from Marguerat et al., 2012. A conservative FDR p-adjusted value of 0.1 yielded 1207 UR and 1483 DR genes (Supplementary Table 12). Of the 973 UR protein-coding genes, 100 were in common with the *hid1Δ* UR genes and were over-represented for molecular function in genes with RNA polymerase II transcription factor activity (Supplementary Table 13). Of the 23 transcription factor genes that were present in our UR set, 19 were in common with those up-regulated in quiescent cells (Table 2). This overlap represented stress or nutrient signalling transcription factors, including the genes encoding Atf1 and Pap1. The modules involving Atf1 and Pap1 transcriptional activity had been identified as over-represented in the AnGeLi EA as well. Over-represented in biological processes were also genes involved in conjugation and cellular fusion and cellular response to starvation. There was also substantial overlap between *hid1Δ* UR genes and quiescence DR genes (Supplementary Table 14), which indicated that certain processes, such as redox metabolism, cytosolic protein translation and nuclear protein import were active in proliferating *hid1Δ*, whereas they were reduced in quiescent cells. There were 81 genes in common for *hid1Δ* DR and the DR genes of the quiescent cells (Supplementary Table 15). However, here was very little indication of DR of processes common to both sets, with the possible exception of small nitrogen containing molecules.

**Table 2.** Transcription Factor genes corresponding to GO:0000981 UR in *hid1Δ*. Determined by Gene Enrichment Analysis of Molecular Function (www.geneontology.org). The FDR p-adjusted value for this category using the Fisher’s Exact test was 0.001. The log_2_-fold values are from the DESeq2 differential expression analysis of the STAR mapping output.

| Name | Log<sub>2</sub>_fold | Name description/Product |
| --- | --- | --- |
| *sfp1* * | 1.13 | Zinc finger protein sfp1 |
| *SPAC2H10.01* * | 1.06 | DNA-binding transcription factor, zf-fungal binuclear cluster type |
| *gsf1* * | 1.02 | Repressor of FLocculation |
| *SPBC530.11c* * | 0.96 | DNA-binding transcription factor, zf-fungal binuclear cluster type |
| *esc1* * | 0.91 | DNA-binding transcription factor Esc1 |
| *atf1* * | 0.91 | G1 Arrest Defective, M Twenty-Six binding protein |
| *loz1* * | 0.89 | Loss Of Zinc sensing |
| *scr1* * | 0.89 | *S. pombe* CREA homolog |
| *pho7* * | 0.81 | PHOSphate metabolism |
| *SPBC56F2.05c* * | 0.76 | DNA-binding transcription factor |
| *gaf1* * | 0.75 | DNA-binding transcription factor Gaf1 |
| *SPCC18.03* * | 0.66 | shuttle craft like transcriptional repressor/ubiquitin-protein ligase E3 |
| *php2* * | 0.66 | CCAAT-binding factor complex subunit Php2 nitrogen starvation |
| *rst2* * | 0.63 | DNA-binding transcription factor Rst2 |
| *moc3* | 0.62 | DNA-binding transcription factor Moc3 |
| *fhl1* * | 0.59 | DNA-binding forkhead transcription factor Fhl1 |
| *SPCC320.03* | 0.56 | DNA-binding transcription factor |
| *fil1* * | 0.56 | DNA-binding transcription factor, zf-GATA type |
| *hsr1* * | 0.56 | Hydrogen peroxide induced Stress Regulator |
| *adn2* | 0.56 | Adhesion defective protein 2 |
| *sak1* * | 0.54 | Suppressor of A Kinase |
| *pap1* * | 0.46 | *S. pombe* AP-1-like gene |
| *SPBC530.08* | 0.40 | DNA-binding transcription factor, membrane-tethered |

\* Common DNA-binding transcription factor activity, RNA polymerase II-specific after Marguerat et al. (2012)

### Proteomics highlights loss of membrane-associated proteins in *hid1Δ*

Since the GA is responsible for the transfer of proteins to various destinations within the cell, it was possible that the levels of individual proteins would also be affected in *hid1Δ*. We employed label-free mass spectrometry on the same cell populations as analysed by transcriptomics. The output from the mass spectrometric analysis comprised 24,703 unique peptides that ultimately yielded 2802 quantifiable proteins of at least 1 unique peptide. Regarding the three Hid paralogs, only Hid1 and Hid2 were detected. Hid1 was present in all samples except the *hid1Δ* mutant, as expected, and Hid2 was detected in at least one sample for each strain, except for BP90. In contrast to transcript levels that were near or above median compared to all other genes, the relative amount of each Hid within the background of all proteins was very low (Supplementary Fig. 11a), which mirrored the study of Marguerat et al., 2012. Differential expression analysis of BP90, *hid1Δ* and *hid3Δ* compared to Cntl gave many fewer differentially abundant proteins than observed for transcripts (Supplementary Tables 16-18). The mutant *hid1Δ* had slightly more proteins increased than decreased in abundance, whereas the opposite was observed for *hid3Δ* (Supplementary Fig. 11b). An EA through PANTHER or AnGeLi using corresponding gene identifiers did not yield statistically significant results for any of the three standard categories, except for DR proteins in *hid3Δ* where mitochondrial processes appeared to be affected predominantly (Supplementary Tables 19-22). However, functional annotation through the DAVID Knowledgebase flagged GA proteins of *hid1Δ* and ER to Golgi transport for *hid3Δ* as two of the few decreased GO terms for the Cellular Compartment and Biological Process classifications, respectively (Supplementary Tables 23-24). Although the statistical analysis did not indicate substantial differential abundance of proteins for *hid1Δ* proteins, the global proteome was altered more for *hid1Δ* as shown by both a PCA (Supplementary Fig. 12) and a correlation analysis of altered ratios in protein abundance ratios when plotted across corresponding transcript ratios (Supplementary Fig. 13). There was an apparent spread in the *hid1Δ*/Cntl protein ratios not observed for those of either BP90/Cntl or *hid3Δ*/Cntl. Since there was an indication through the DAVID functional annotation that levels of certain GA proteins may be altered in *hid1Δ*, we looked at those proteins that were not detected for this strain, but were present in the other strains (Table 3, Supplementary Table 25). Most missing proteins were GA, membrane-associated or other proteins that could be altered in amount if GA function was compromised, such as the Arf and Ric1 GEFs, a mannosyl-transferase, and proteins of the cell-division site or plasma membrane. There were a number of proteins not detected for *hid3Δ* only and these corresponded to membrane proteins of cell organelles, including peroxisomes, endoplasmic reticulum and mitochondria (Supplementary Table 26).

**Table 3.** Proteins missing in the proteome of *hid1Δ*. The proteins listed were missing in all three replicates of *hid1Δ*, but present in all other strains. The Uniprot identifiers and other metadata for each protein are given in Additional file 6.

| Systematic_ID | Description/<br>Product | Cellular<br>Component* |
| --- | --- | --- |
| SPAC607.06c | metallopeptidase | cell division site/cytosol/nucleus |
| SPCC417.05c | SEL1/TPR repeat protein CFH2 | cell division site/cytosol/nucleus |
| SPAC4D7.01c | Arf GEF Sec71 | Golgi Apparatus/Golgi associated vesicle |
| SPCC757.15 | cytochrome c oxidase assembly<br>protein Cox14 | Integral component of membrane |
| SPAC22G7.02 | karyopherin/importin beta family<br>nuclear import signal receptor Kap111 | cytosol/nuclear membrane/nuclear periphery |
| SPBC460.01c | amino acid transmembrane<br>transporter | Integral component of membrane |
| SPAC12G12.08 | mitochondrial ribosomal protein<br>subunit L6 | mitochondrial large ribosomal subunit |
| SPAC17C9.16c | plasma membrane spermidine<br>transmembrane transporter Mfs1 | endoplasmic reticulum/Integral<br>component of membrane/plasma membrane |
| SPBC3H7.11 | tRNA(Ser) residue 32 (cytosine-3-)<br>-methyltransferase Trm141 | cytosol/nucleus |
| SPCC965.13 | plasma membrane pyridoxal family<br>Transmembrane transporter | cell division site/cell tip/endoplasmic reticulum/<br>Golgi apparatus/integral component of plasma<br>membrane |
| SPCC70.08c | methyltransferase | cytosol/nucleus |
| SPCC1919.02 | pig-X, glycosylphosphatidyl-inositol-<br>mannosyltransferase complex,<br>subunit I | endoplasmic reticulum membrane/integral<br>component of membrane |
| SPAC17A5.16 | human HID1 ortholog 3 | Ftp105 (Hid1) Golgi apparatus |
| SPAC1851.04c | Rab-specific(Ryh1/Ypt6) Ric1-Rgp1<br>guanyl-nucleotide exchange factor,<br>subunit Ric1/Sat4 | cytosol/Golgi membrane/integral component<br>of membrane |
| SPBC359.05 | vacuolar heme ABC transmembrane<br>exporter Abc3 | fungus-type vacuole membrane/<br>integral component of membrane |
| SPAC222.04c | Ino80 complex subunit Ies6 | chromatin/cytoplasm/Ino80 complex |
| SPBC27B12.09c | mitochondrial carrier FAD Flx1 | integral component of membrane/<br>mitochondrial inner membrane |
| SPAC644.04 | RNA 5'-triphosphatase Pct1 | mRNA cap methyltransferase complex |

\* Putative Cellular Component was taken from Pombase (www.pombase.org)

### *hid1Δ* grows poorly on minimal medium

Initial efforts to create replacement mutants using standard auxotrophic complementation yielded few colonies for *hid1* and resulted mainly in false positives. We surmised that the use of minimal medium for heterotrophic selection was problematic. We tested this possibility using standard drop tests of decreasing cell amounts on agar plates to compare proliferation on rich YE medium and Edinburgh Minimal Medium. Although all strains exhibited reduced colony growth with EMM compared to YE, *hid1Δ* was particularly affected on the minimal medium with even high-density spots barely being visible (Fig. 4). The discrepancy between rich and minimal media suggests that some nutrient or component from the minimal medium may be limiting within cells of *hid1Δ* to support its proliferation.

**Figure 4.**
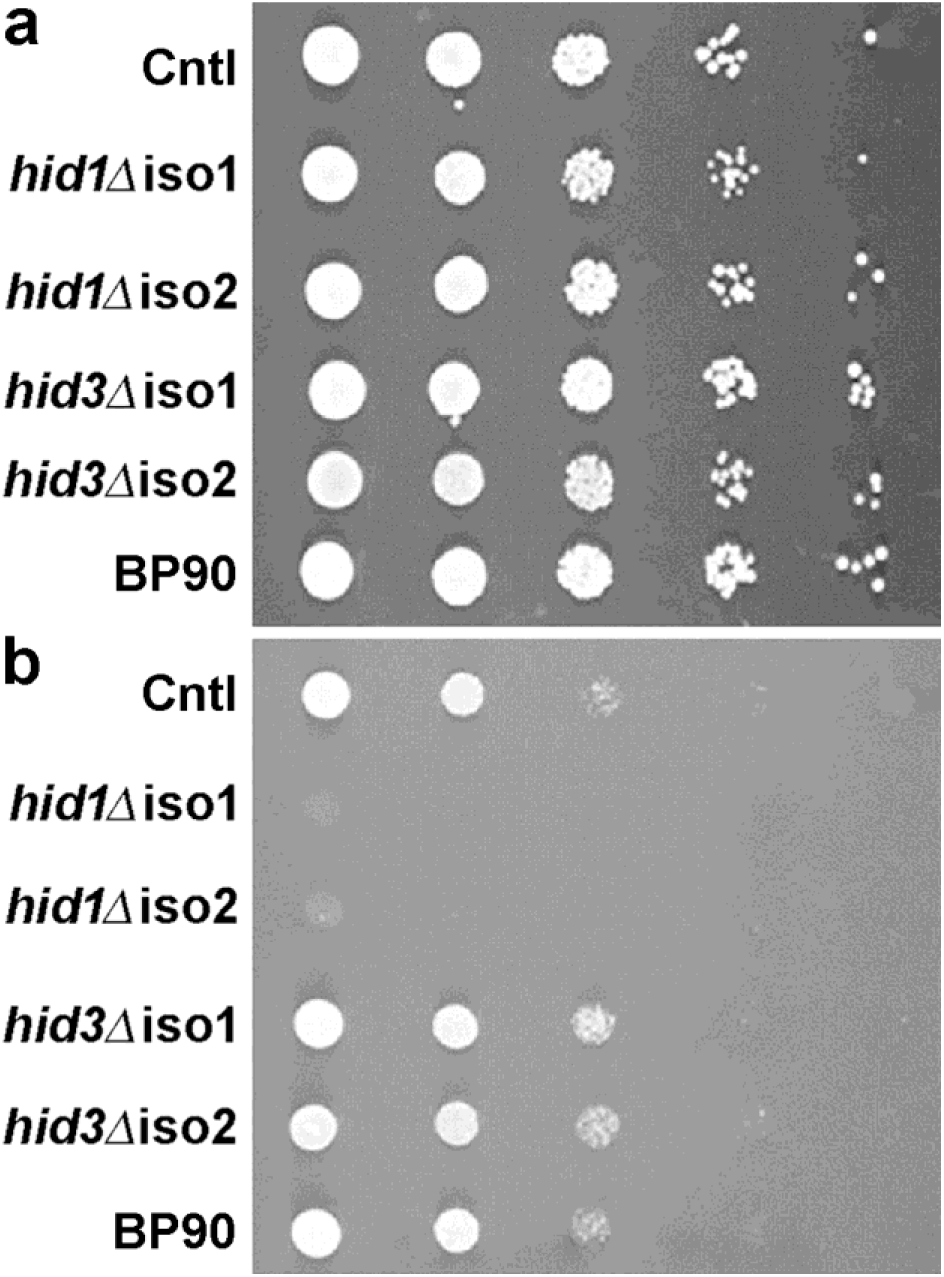
Differential proliferation of mutants on minimal medium. Cells of each mutant strain were grown overnight in standard YES at 30 °C with shaking at 200 RPM. The cells were counted using the Cellometer™ Mini (Ozyme Ltd) and a series of 10-fold dilutions of cells were made starting from 5×10^6^ cells/ml (left column). A volume of 5 μl was placed for each spot onto petri dishes containing either YES/agar (A) or Edinburgh Minimal Medium (B) plus supplements. EMM was purchased from Formedium Ltd (Norfolk, United Kingdom). Plates were incubated for 4 days before images were taken.

## Discussion

The suggestion that human HID1 was a potential tumor suppressor and the subsequent isolation of the *C. elegans* ortholog as a potential repressor of the Dauer state raised a potential contradiction in functional interpretation of HID1 as a factor restricting or maintaining cell proliferation. A direct approach to address this apparent contradiction was to evaluate *HID1* expression in cancers and explore inherent bias that might have been due to use of cultured cancer cells (Harada et al. 2001). Although gene expression between cell lines and directly obtained patient-derived tumors appears to be positively correlated in general (Ghandi et al. 2019), this may not hold for individual genes. Our approach to relate cell line versus primary tumor expression for individual genes revealed that cell-line expression analysis does not serve as an appropriate model for expression analysis of *HID1* in most cancer types. The inability of Celligner to correct *HID1* expression between cell lines and tumors supported our finding that apparent *HID1* expression is lower in cell lines. *HID1* appears to be infrequently mutated, but frequently amplified, in tumors compared to other GA-related genes. These observations are inconsistent with a reduced expression accelerating cell proliferation. Any mutation or decrease in expression leading to less HID1 would result more likely in reduced proliferation. Some genes were observed to have either systematic or cancer-specific differences in transcript levels among cell lines and tumors indicating that a cell line(s) must be verified on an individual gene basis before being employed as a model for expression in tumors. In addition to *HID1*, our approach identified the GA genes *DYM* and the *JANUS KINASE AND MICROTUBULE INTERACTING PROTEINs (JAKMIP/NECC)* whose transcript levels appeared to be reduced in cell lines. A possible connection among the genes is structural maintenance of the GA. The GA has a modified structure in cells lacking DYM (Dimitrov et al. 2009) and pharmaceutical treatment or mutation that disrupts microtubule interaction causes GA dissociation (Ayscough et al. 1993; Chabin-Brion et al. 2001). We have recently reported that elimination of Hid1 in *S. pombe* appears to remove GA stacking capability (Hooks et al. 2025).

A direct time-course analysis of cell counts in liquid culture of the various mutant strains confirmed previous observations of the slow proliferation phenotype of *hid1Δ* (Hooks et al. 2025). This reduced proliferation is attributed to an increase in the lag-time to recover a normal rate of proliferation upon re-culturing. We speculated that the mutant was less capable than other strains to recovery from the metabolic changes initiated under stationary phase. The indication that transport capacity may be perturbed in *hid1Δ* comes from our holistic profiling, particularly of transcriptional responses. The major UR genes encode cell surface proteins and transporters. Unlike *E. nidulans* (Dimou et al. 2020), plasma membrane transporters are trafficked through the GA in *S. pombe* (Nakase et al. 2012). For example, two of the top UR genes encode flocculin-like proteins that are cell surface proteins involved in cell adhesion (Matsuzawa et al. 2011) and response to environmental changes (Bouyx et al. 2021). Elimination of Hid1 appears to affect cell conjugation as revealed through a systematic screen to identify factors important for meiosis in *S. pombe* (Blyth et al. 2018). Furthermore, the UR of cell adhesion genes appears to be a common consequence to the onset of quiescence (for review, see Marescal and Cheeseman, 2020). Thus, the UR of genes encoding metabolite and ion transporters suggests a metabolic deficiency for *hid1Δ*. The contribution of GA function to maintaining cellular quiescence has been reported, whereby GA associated proteins reduced cell viability under quiescence imposed by nitrogen-starvation (Sajiki et al. 2009). This study also reported an important UR of RNA polymerase II processes to cellular quiescence, a feature that we observed in our transcriptomic profiling. The induction of gene expression requires the activation of signalling pathways and subsequent transcription factor activity. We observed a substantial overlap of TF genes induced in *hid1Δ* with those of nitrogen-starved cells (Marguerat et al. 2012). Nevertheless, since *hid1Δ* is not quiescent at the point of analysis, differences between the Hid1 deficiency and nutritionally imposed quiescence would be expected. The overlapping TF genes were related to both stress and nutrition, whereas the few UR TF genes unique to our study pertain to cell surface or cell exterior processes. A systematic screen of *S. pombe* mutant strains for alteration in chronological lifespan during nutrient-related quiescence noted a reduction in viability for *hid1Δ* (Romila et al. 2021). However, it was noted that genetic contributions to lifespan are context-dependent to the genetic background and the method of monitoring viability or proliferation. It is possible that the apparent reduction in viability was due to an increase in the lag-time that subsequently reduced the barcode profile of *hid1Δ* in this screen. Any reduction in metabolite transport capacity resulting from starvation-induced quiescence may not recover as quickly upon return to favourable growth conditions with GA function, and thus protein trafficking, being compromised.

More and more, examples are forthcoming showing a relationship between the perturbation of GA function and negative effects on cell function and organism physiology, from tumorigenesis and neurodegenerative diseases to severe genetic abnormalities (Machamer, 2015). There is a link between cellular quiescence and a delay in tumorigenesis (Torrano and Carracedo, 2017), therefore, a variety of models for studying cellular and metabolic quiescence will be useful, including those that do not require a dramatic modification of nutrient regimes (Gray et al. 2004; Sagot and Laporte, 2019). Even for eukaryotic microorganisms, which host a large variety of GA structures and appear physiologically robust to the loss of GA proteins and structure, disruption of GA function affects gene expression and may have more profound consequences depending on cellular metabolic state. Thus, models like the *S. pombe hid1Δ* mutant, may be informative as to metabolic processes that occur during the lag-phase or the recovery from lag-phase, and which have applications in various biological processes from health to food science (Bertrand, 2019).

## Acknowledgements

We thank those at the Centre Genomique Fonctionelle de Bordeaux, particularly Dr. Christophe Hubert at the Sequencing facility and Dr. Macha Nicolski and Dr. Aurelien Barré at the Centre de Bioinformatique. The results shown here are in part based upon data generated by the TCGA Research Network and the Cancer Cell Line Encyclopedia.

## Author information

### Contributions

A-PF created and characterized mutant strains, mined the transcriptomic data sets and helped write the manuscript. AA contributed the phylogenetic analysis. MA created and characterized mutant strains. KBH conducted the analysis of CCLE and TCGA data sets. LB conducted the TEM. SC collected and analysed the proteomic data and HF validated and conducted subsequent database interrogations. RFM helped plan the experimental program and contributed to manuscript preparation. MAH was the principal investigator and the corresponding author of the study.

### Present addresses

A-PF: Firmenich SA, Rue de Bergere 7, 1242 Satingy Geneva, Switzerland

AA, MA: Faculty of Science, University of Tabuk, Kingdom of Saudi Arabia

KBH: Selvita S.A. Podole 7930-394 Krakow, Poland

### Data Availability

The data sets to support the work will be released upon publication.

## References

Ailion M, Thomas JH. 2003. Isolation and characterization of high-temperature-induced Dauer formation mutants in Caenorhabditis elegans. Genetics 165: 127–144 doi:10.1093/genetics/165.1.127

Ayala I, Colanzi A. 2017. Alterations of golgi organization in Alzheimer’s disease: a cause or a consequence? Tissue Cell 49: 133–140. doi:10.1016/j.tice.2016.11.007

Aydin B, Arga KY. 2019. Co-expression network analysis elucidated a core module in association with prognosis of non-functioning non-invasive human pituitary adenoma. Front Endocrinol 10: 361. doi:10.3389/fendo.2019.00361

Ayscough K, Hajibagheri NMA, Watson R and Warren G. 1993. Stacking of Golgi cisternae in Schizosaccharomyces pombe requires intact microtubules. J Cell Sci 106: 1227–1237. doi:10.1242/jcs.106.4.1227

Bähler J, Wu JQ, Longtine MS, Shah NG, McKenzie A, Steever AB, Wach A, Philippsen P, Pringle JR. 1998. Heterologous modules for efficient and versatile PCR-based gene targeting in *Schizosaccharomyces pombe*. Yeast 14: 943–951. doi:10.1002/(SICI)1097-0061(199807)14:10

Barlow LD, Nývltová E, Aguilar M, Tachezy J, Dacks JB. 2018. A sophisticated, differentiated golgi in the ancestor of eukaryotes. BMC Biol 16: 27. doi:10.1186/s12915-018-0492-9

Bernard P, Hardwick K, Javerzat JP. 1998. Fission yeast bub1 is a mitotic centromere protein essential for the spindle checkpoint and the preservation of correct ploidy through mitosis. J. Cell Biol. 143: 1775–1787. 10.1083/JCB.143.7.1775.

Bertrand RL. 2019. Lag Phase Is a dynamic, organized, adaptive, and evolvable period that prepares bacteria for cell division. J Bacteriol 201: e00697–18. doi:10.1128/JB.00697-18

Blyth J, Makrantoni V, Barton RE, Spanos C, Rappsilber J, Marston AL. 2018. Genes important for schizosaccharomyces pombe meiosis identified through a functional genomics screen. Genetics 208: 589–603. doi:10.1534/genetics.117.300527

Bouyx C, Schiavone M, François JM. 2021. FLO11, a developmental gene conferring impressive adaptive plasticity to the yeast *Saccharomyces cerevisiae*. Pathogens 10: 1509. doi:10.3390/pathogens10111509

Chabin-Brion K, Marceiller J, Perez F, Settegrana C, Drechou A, Durand G, Poüs C. 2001. The Golgi Complex Is a Microtubule-organizing Organelle. Mol Biol Cell 12: 2047. doi:10.1091/mbc.12.7.2047

Dimitrov A, Paupe V, Gueudry C, Sibarita J-B, Raposo G, Vielemeyer O, Gilbert T, Csaba Z, Attie-Bitach T, Cormier-Daire V, et al. 2009. The gene responsible for Dyggve-Melchior-Clausen syndrome encodes a novel peripheral membrane protein dynamically associated with the Golgi apparatus. Hum Mol Genet 18: 440–53. doi:10.1093/hmg/ddn371

Dimou S, Martzoukou O, Dionysopoulou M, Bouris V, Amillis S, Diallinas G. 2020. Translocation of nutrient transporters to cell membrane via Golgi bypass in Aspergillus nidulans. EMBO Rep 21: e49929. doi:10.15252/embr.201949929

Du W, Zhou M, Zhao W, Cheng D, Wang L, Lu J, Song E, Feng W, Xue Y, Xu P, et al. 2016. HID-1 is required for homotypic fusion of immature secretory granules during maturation. ELife 5: e18134. doi:10.7554/eLife.18134

Dupuis N, Lebon S, Kumar M, Drunat S, Graul-Neumann LM, Gressens P, El Ghouzzi V. 2013. A novel RAB33B mutation in Smith-McCort dysplasia. Hum Mutat 34: 283–286. doi:10.1002/humu.22235

El Ghouzzi V, Dagoneau N, Kinning E, Thauvin-Robinet C, Al-Gazali WC, Prost-Squarcioni C, Lihadh I, Verloes A, Merrer ML, Munnich A, et al. 2003. Mutations in a novel gene DYMeclin (FLJ20071) are responsible for Dyggve-Melchior-Clausen syndrome. Hum Mol Genet 12: 357–364. doi:10.1093/hmg/ddg029

Feichtinger J, Aldeailej I, Anderson R, Almatrafi M, Alsiwiehri N, Griffiths K, Stuart N, Wakeman JA, Larcombe L, McFarlane RJ. 2012. Meta-analysis of clinical data using human meiotic genes identifies a novel cohort of highly restricted cancer-specific marker genes. Oncotarget 3: 843–853. doi:10.18632/oncotarget.580

Feng Y, He F, Yan S, Huang H, Huang Q, Deng T, Wu H, Gao B, Liu J. 2017. The Role of GOLPH3L in the Prognosis and NACT response in Cervical Cancer. J Cancer 8: 443–454. doi:10.7150/jca.17096

Ghandi M, Huang FW, Jané-Valbuena J, Kryukov G V., Lo CC, McDonald ER, Barretina J, Gelfand ET, Bielski CM, Li H, et al (2019) Next-generation characterization of the Cancer Cell Line Encyclopedia. Nature 569: 503–508. doi:10.1038/s41586-019-1186-3

Gray J V, Petsko GA, Johnston GC, Ringe D, Singer RA, Werner-Washburne M. 2004. ‘Sleeping beauty’: quiescence in Saccharomyces cerevisiae. Microbiol Mol Biol Rev 68: 187–206. doi:10.1128/MMBR.68.2.187-206.2004

Harada H, Nagai H, Tsuneizumi M. 2001. Identification of DMC1, a novel gene in the TOC region on 17q25.1 that shows loss of expression in multiple human cancers. J Hum Genet 46: 90–95. doi:10.1007/s100380170115

Hooks MA, Alasmari A, Alshehri M, Brocard L, Hooks KB, Julien M, Mcfarlane RJ. 2025. Golgi_traff phylogeny reveals ancient eukaryotic genes with recent surprises: replication and diversification of HID1 domain-containing protein unique to Schizosaccharomyces. FEMS Microbiol. Lett. 372: FNAF088. 10.1093/FEMSLE/FNAF088.

Hu ZY, Liu YP, Xie LY, Wang XY, Yang F, Chen S-Y, Li ZG. 2016. AKAP-9 promotes colorectal cancer development by regulating Cdc42 interacting protein 4 Biochim Biophys Acta 6: 1172–1181. doi:10.1016/j.bbadis.2016.03.012

Hummer BH, de Leeuw NF, Burns C, Chen L, Joens MS, Hosford B, Fitzpatrick JAJ, Asensio CS. 2017. HID-1 controls formation of large dense core vesicles by influencing cargo sorting and trans-Golgi network acidification. Mol Biol Cell 28: 3870–3880. doi:10.1091/mbc.E17-08-0491

Hummer B, Carter T, Sellers B, Triplett J, Asensio C. 2023. Identification of the functional domain of the dense core vesicle biogenesis factor HID-1. PLoS ONE 9: e0291977. 10.1371/journal.pone.0291977.

Johnson A, Severson E, Gay L, Vergilio J-A, Elvin J, Suh J, Daniel S, Covert M, Frampton GM, Hsu S, et al. 2017. Comprehensive Genomic Profiling of 282 Pediatric Low- and High-Grade Gliomas Reveals Genomic Drivers, Tumor Mutational Burden, and Hypermutation Signatures. Oncologist 22: 1478–1490. doi:10.1634/theoncologist.2017-0242

Kennedy PJ, Vashisht AA, Hoe KL, Kim DU, Park HO, Hayles J, Russell P. .2008. A genome-wide screen of genes involved in cadmium tolerance in Schizosaccharomyces pombe. Toxicol. Sci. 106: 124–139. 10.1093/TOXSCI/KFN153.

Kim DU, Hayles J, Kim D, Wood V, Park H, Won M, Yoo H, Duhig T, Nam M, Palmer G, et al. 2010. Analysis of a genome-wide set of gene deletions in the fission yeast Schizosaccharomyces pombe. Nature biotechnology 28: 617–23. 10.1038/nbt.1628.

Kouranti I, McLean J, Feoktistova AA. 2010. A global census of fission yeast deubiquitinating enzyme localization and interaction networks reveals distinct compartmentalization profiles and overlapping functions in endocytosis and polarity. PLoS Biol 8: 9 doi:10.1371/journal.pbio.1000471

Longo T, Mcginley KF, Freedman JA, Etienne W, Wu Y, Sibley A, Owzar K, Gresham J, Moy C, Szabo S, et al. 2016. Targeted Exome Sequencing of the Cancer Genome in Patients with Very High-risk Bladder Cancer. Eur Urol 70: 714–717. doi:10.1016/j.eururo.2016.07.049

Lyne R, Burns G, Mata J, Penkett CJ, Rustici G, Chen D, Langford C, Vetrie D, Bähler J. 2003. Whole-genome microarrays of fission yeast: Characteristics, accuracy, reproducibility, and processing of array data. BMC Genomics 4: 1–15. doi:10.1186/1471-2164-4-27

Machamer CE. 2015. The Golgi complex in stress and death. Front Neurosci 9: 421 doi:10.3389/fnins.2015.00421

Makowski SL, Tran TT, Field SJ. 2017. Emerging themes of regulation at the Golgi. Curr Opin Cell Biol 45: 17–23. doi:10.1016/j.ceb.2017.01.004

Marescal O, Cheeseman IM. 2020. Cellular mechanisms and regulation of quiescence. Dev Cell 55: 259–271. doi:10.1016/j.devcel.2020.09.029

Marguerat S, Schmidt A, Codlin S, Chen W, Aebersold R, Bähler J. 2012. Quantitative analysis of fission yeast transcriptomes and proteomes in proliferating and quiescent cells. Cell 151: 671–83. doi:10.1016/j.cell.2012.09.019

Matsuyama A, Shirai A, Yoshida M. 2008. A series of promoters for constitutive expression of heterologous genes in fission yeast. Yeast 25: 371–6. doi:10.1002/yea.1593

Matsuzawa T, Morita T, Tanaka N, Tohda H, Takegawa K. 2011. Identification of a galactose-specific flocculin essential for non-sexual flocculation and filamentous growth in *Schizosaccharomyces pombe*. Mol Microbiol 82: 1531–1544. doi:10.1111/j.1365-2958.2011.07908.x

Mayinger P. 2011. Signaling at the golgi. Cold Spring Harb Perspect Biol 3: 1–14. doi:10.1101/cshperspect.a005314

McFarlane RJ, Wakeman JA. 2017 Meiosis-like functions in oncogenesis: A new view of cancer. Cancer Res 77: 5712–5716. doi:10.1158/0008-5472.CAN-17-1535

Mesa R, Luo S, Hoover CM, Miller K, Minniti A, Inestrosa N, Nonet ML. 2011. HID-1, a new component of the peptidergic signaling pathway. Genetics 187: 467–483. doi:10.1534/genetics.110.121996

Nakase M, Nakase Y, Chardwiriyapreecha S, Kakinuma Y, Matsumoto T, Takegawa K. 2012. Intracellular trafficking and ubiquitination of the Schizosaccharomyces pombe amino acid permease Aat1p. Microbiology 158: 659–673. doi:10.1099/mic.0.053389-0

Popławski P, Piekiełko-Witkowska A, Nauman A. 2017. The significance of TRIP11 and T3 signalling pathway in renal cancer progression and survival of patients. Endokrynol Pol 68: 631–641. doi:10.5603/EP.a2017.0052

Rodríguez-López M, Bordin N, Lees J, Scholes H, Hassan S, Saintain Q, Kamrad S, Orengo C, Bähler J, et al. 2023. Broad functional profiling of fission yeast proteins using phenomics and machine learning. Elife 12: RP88229–RP88229. 10.7554/ELIFE.88229.

Romila CA, Townsend SJ, Malecki M, Kamrad S, Rodríguez-López M, Hillson O, Cotobal C, Ralser M, Bähler J. 2021. Barcode sequencing and a high-throughput assay for chronological lifespan uncover ageing-associated genes in fission yeast. Microb Cell 8: 146–160. doi:10.15698/mic2021.07.754

Sagot I, Laporte D. 2019. The cell biology of quiescent yeast - a diversity of individual scenarios. J Cell Sci 132: 1–10. doi:10.1242/jcs.213025

Sajiki K, Hatanaka M, Nakamura T, Takeda K, Shimanuki M, Yoshida T, Hanyu Y, Hayashi T, Nakaseko Y, Yanagida M. 2009. Genetic control of cellular quiescence in *S. pombe*. J Cell Sci 122: 1418–1429. doi:10.1242/jcs.046466

Sechi S, Frappaolo A, Karimpour-Ghahnavieh A, Piergentili R, Giansanti MG. 2020. Oncogenic roles of GOLPH3 in the physiopathology of cancer. Int J Mol Sci 21: 933. doi:10.3390/ijms21030933

Schänzer A, Achleitner MT, Trümbach D, Hubert L, Munnich A, Ahlemeyer B, AlAbdulrahim M, Greif PA, Vosberg S, Hummer B, et al. 2021. Mutations in HID1 Cause Syndromic Infantile Encephalopathy and Hypopituitarism. Ann. neurol. 90: 143–158 10.1002/ANA.26127.

Sharabi K, Charar C, Friedman N, Mizrahi I, Zaslaver A, Sznajder JI, Gruenbaum Y. 2014. The Response to High CO2 Levels Requires the Neuropeptide Secretion Component HID-1 to Promote Pumping Inhibition. PLoS Genetics 10: e1004529. doi:10.1371/journal.pgen.1004529

Soneson C, Robinson MD. 2018. Bias, robustness and scalability in single-cell differential expression analysis. Nat Meth 15: 255–261. doi:10.1038/nmeth.4612

Torrano V, Carracedo A. 2017. Quiescence-like metabolism to push cancer out of the race. Cell Metab 25: 997–999. doi:10.1016/j.cmet.2017.04.027

Wang L, Zhan Y, Song E, Yu Y, Jiu Y, Du W, Lu J, Liu P, Xu P, Xu T. 2011. HID-1 is a peripheral membrane protein primarily associated with the medial- and trans-Golgi apparatus. Protein Cell 2: 74. doi:10.1007/s13238-011-1008-3

Warren A, Chen Y, Jones A, Shibue T, Hahn WC, Boehm JS, Vazquez F, Tsherniak A, McFarland JM. 2021. Global computational alignment of tumor and cell line transcriptional profiles. Nat Commun 12: 1–12. doi:10.1038/s41467-020-20294-x

Wood V, Gwilliam R, Rajandream M-A, Lyne M, Lyne R, Stewart A, Sgouros J, Peat N, Hayles J, Baker S, et al. 2002. The genome sequence of Schizosaccharomyces pombe. Nature 415: 871–80 doi:10.1038/nature724

Yu Y, Wang L, Jiu Y, Zhan Y, Liu L, Xia Z, Song E, Xu P, Xu T. 2011. HID-1 is a novel player in the regulation of neuropeptide sorting. Biochem J 434: 383–390. doi:10.1042/BJ20110027

