## Supplementary Figures for "HID1 domain-containing protein 1 is required for normal cell proliferation in *Schizosaccharomyces pombe*"

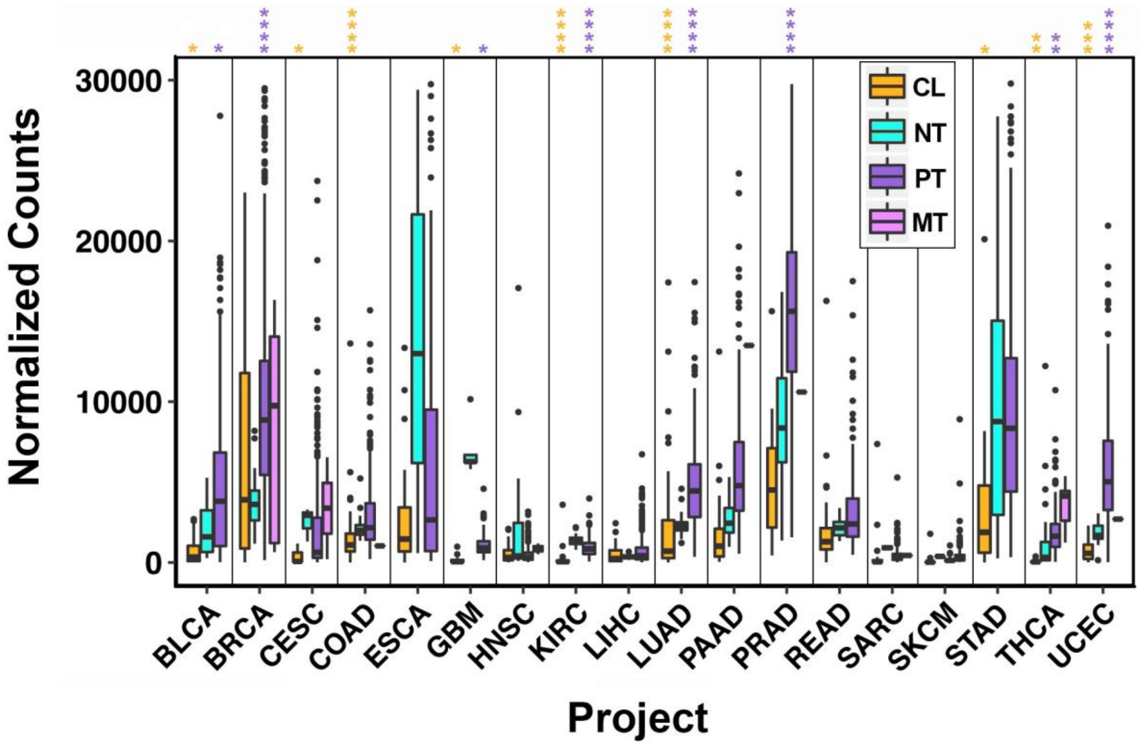


**Supplementary Fig. 1.** Boxplot presentation of *HID-1* transcript counts in sample types representing various cancers. RNAseq-based transcript counts were obtained from TCGA projects having both normal tissue and primary tumor data. RNAseq-based cell-line transcript counts were obtained from the Cancer Cell-line Encyclopedia and matched to TCGA projects. The relative expression values were normalized using DeSEQ2 for each project independently. Legend: CL, cell line; NT, normal tissue, PT, primary tumor; MT, metastatic tumor (where available). The black dots represent sample outliers. The stars represent the statistical significance according to Mood's median test with an FDR (BH) = 0.05: *, $p < 0.05$; ** $p < 0.01$; *** $p < 0.001$; ****, $p < 0.0001$.

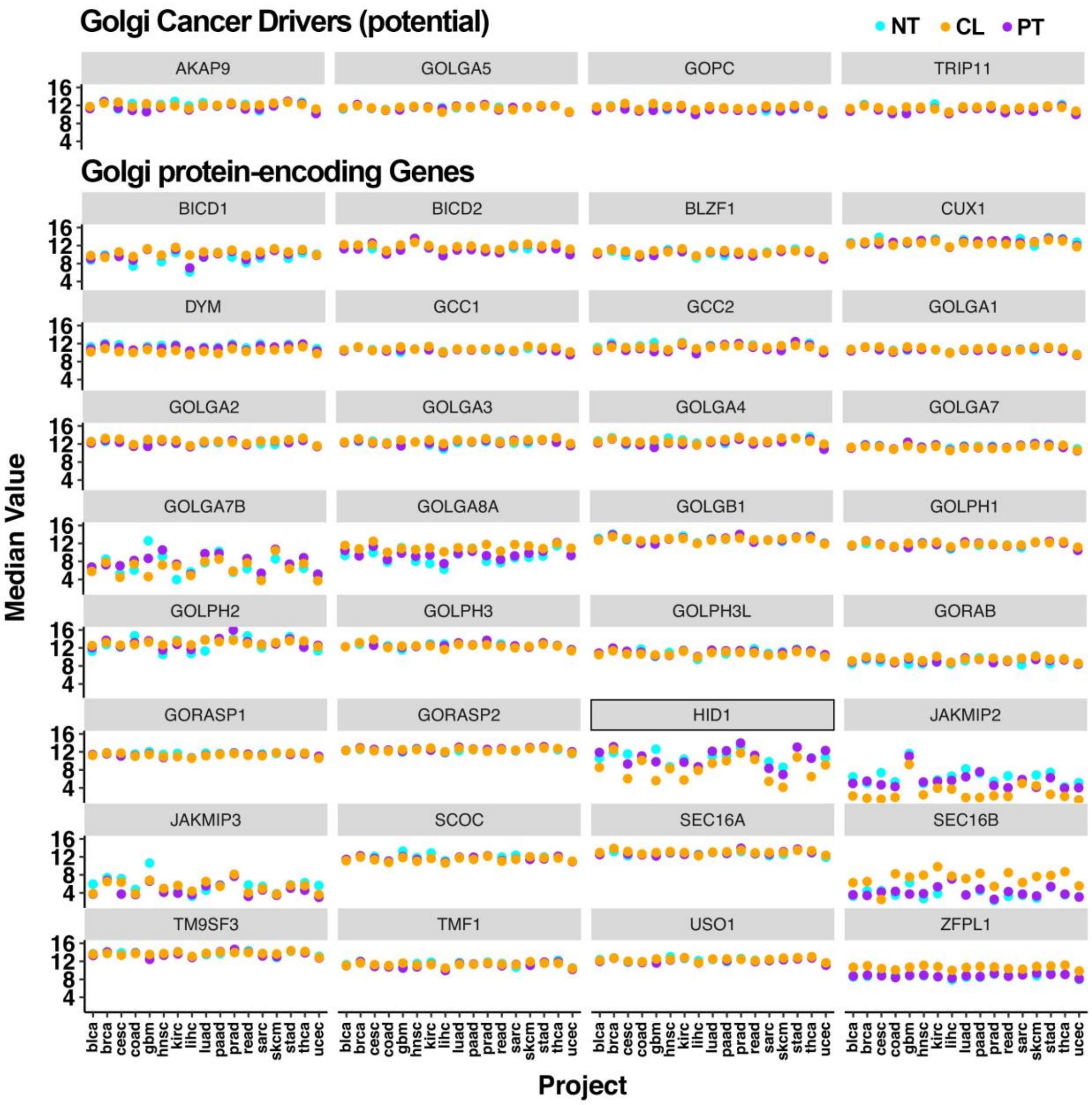
Golgi Cancer Drivers (potential)
NT CL PT
AKAP9
GOLGA5
GOPC
TRIP11
Golgi protein-encoding Genes
BICD1
BICD2
BLZF1
CUX1
DYM
GCC1
GCC2
GOLGA1
GOLGA2
GOLGA3
GOLGA4
GOLGA7
GOLGA7B
GOLGA8A
GOLGB1
GOLPH1
GOLPH2
GOLPH3
GOLPH3L
GORAB
GORASP1
GORASP2
HID1
JAKMIP2
JAKMIP3
SCOC
SEC16A
SEC16B
TM9SF3
TMF1
USO1
ZFPL1
Median Value
16
12
8
4
blca brca cesc coad gbm hnsc kirc lihc luad paad prad read sarc skcm stad thca ucec
Project

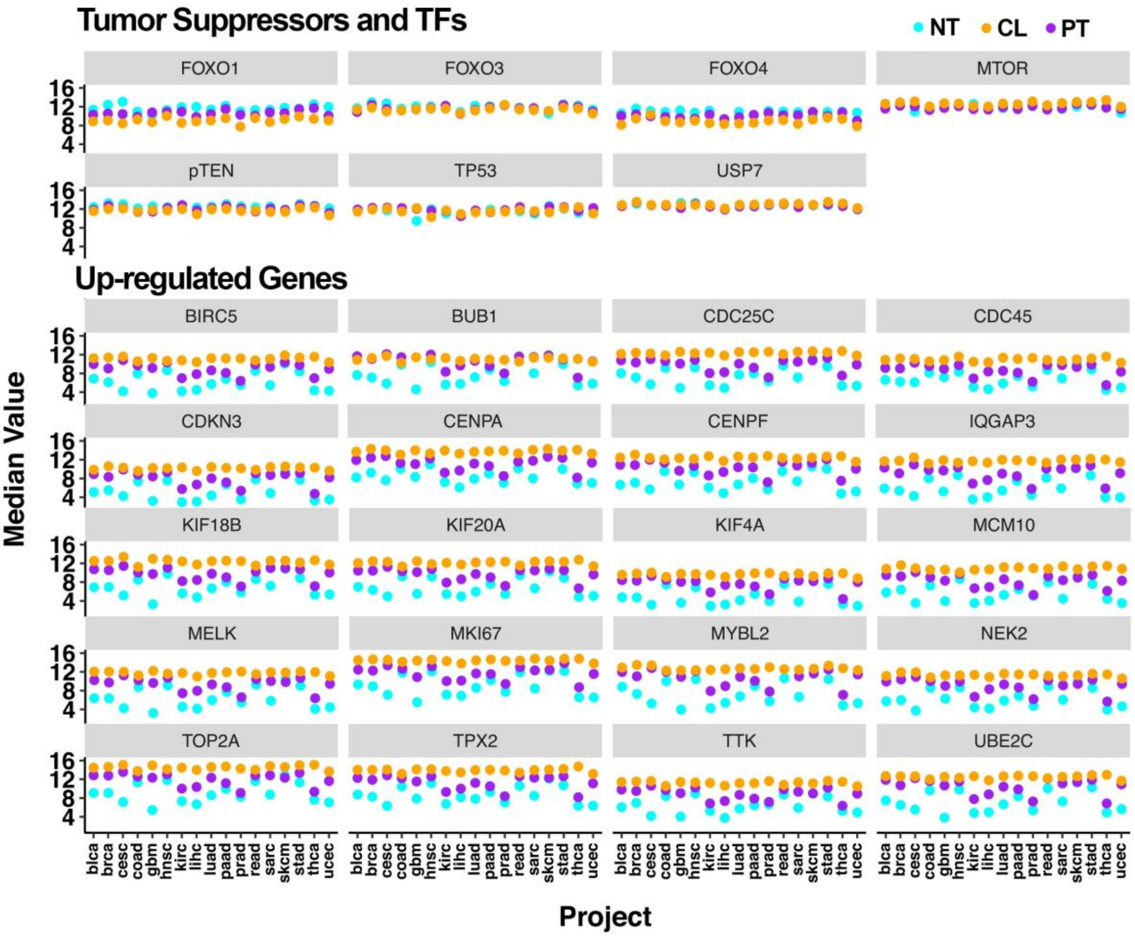


**Supplementary Fig. 2.** Comparison of transcript counts among normal tissues, primary tumors and cell lines across various cancers. Each dot represents the median value of normalized transcript counts for samples in a respective cancer type according to the TCGA project designation. The black box highlights *HID1,* which is the dot representation of the data in Supplementary Figure 1. Abbreviations: N, normal tissue; P, primary tumor; C, cancer cell line.

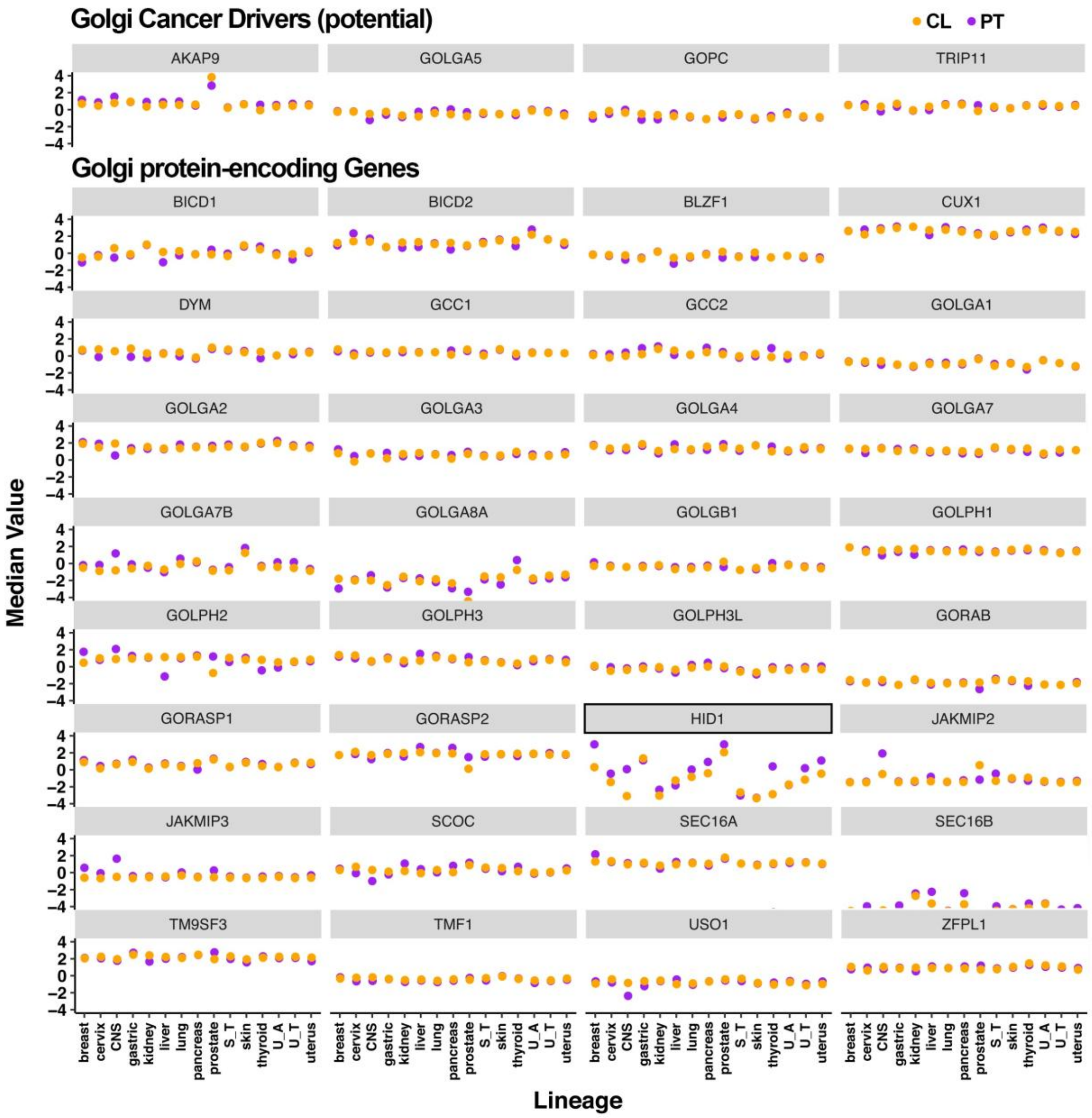

Golgi Cancer Drivers (potential)
CL
PT
AKAP9
GOLGA5
GOPC
TRIP11
Golgi protein-encoding Genes
BICD1
BICD2
BLZF1
CUX1
DYM
GCC1
GCC2
GOLGA1
GOLGA2
GOLGA3
GOLGA4
GOLGA7
GOLGA7B
GOLGA8A
GOLGB1
GOLPH1
GOLPH2
GOLPH3
GOLPH3L
GORAB
GORASP1
GORASP2
HID1
JAKMIP2
JAKMIP3
SCOC
SEC16A
SEC16B
TM9SF3
TMF1
USO1
ZFPL1
Median Value
4
2
0
−2
−4
breast
cervix
CNS
gastric
kidney
liver
lung
pancreas
prostate
S_T
skin
thyroid
U_A
U_T
uterus
Lineage

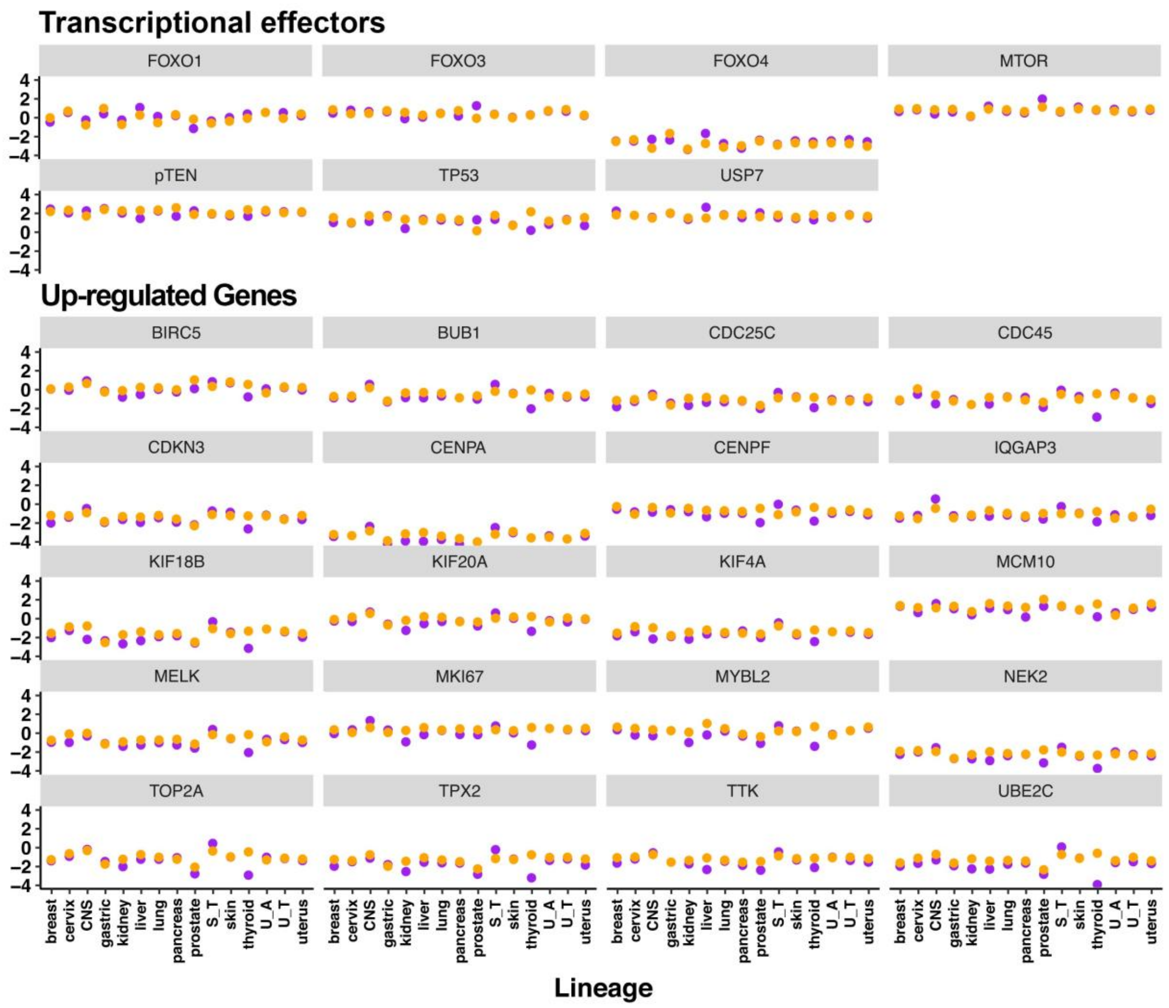


**Supplementary Fig. 3.** Comparison of Celligner-normalized transcript counts between primary tumors and cell lines. Each dot represents the median value of normalized transcript counts for samples of cancer types according to the tissue of origin. The black box highlights *HID1.* Abbreviations: CL, cancer cell line; PT, primary tumor.

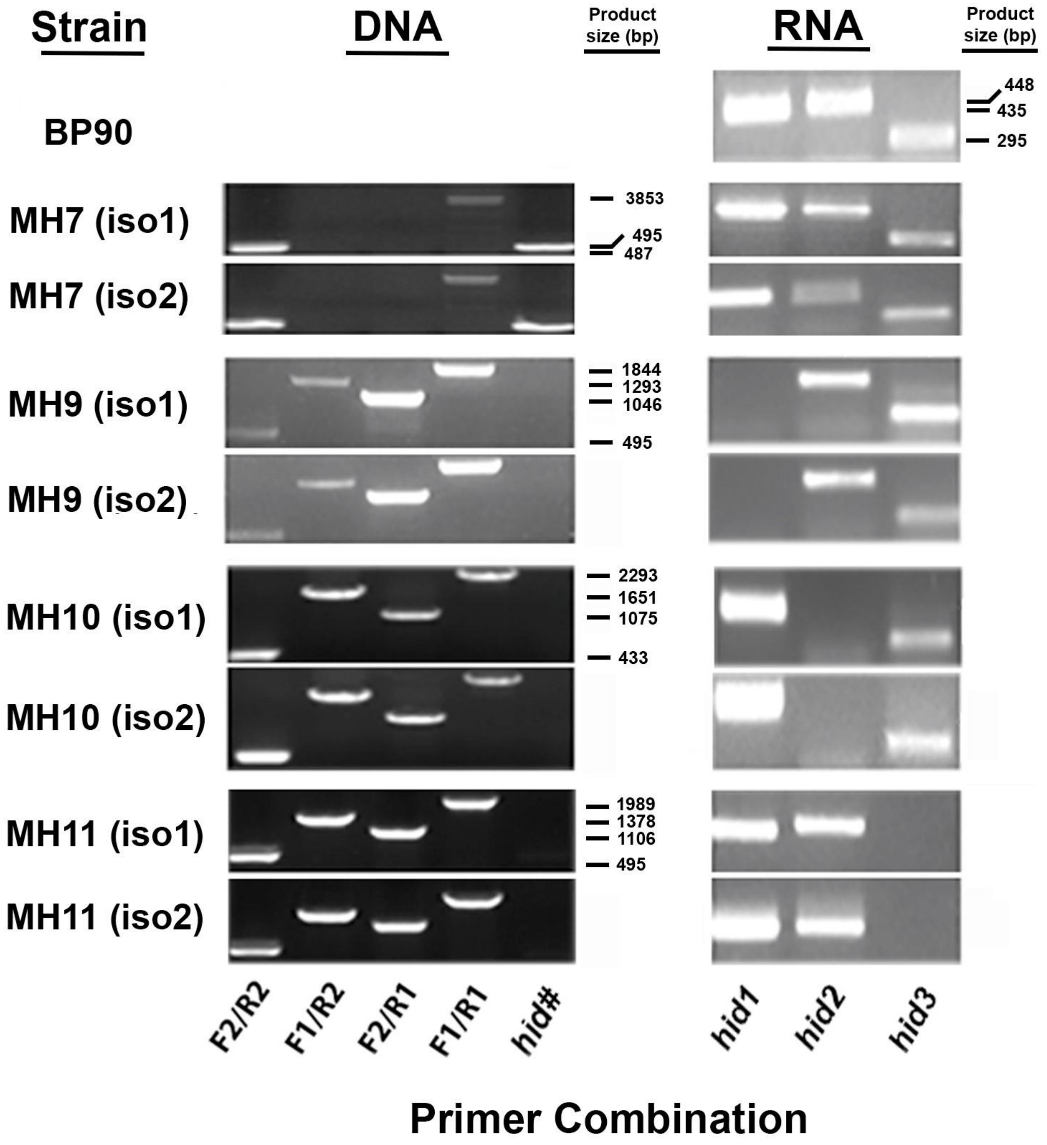


**Supplementary Fig. 4.** Confirmation of gene replacement and *hid* expression in mutants as determined by PCR and RT-PCR. All replacement mutant strains were given number designations according to the order they were selected from plates for subsequent genotyping. DNA and total RNA was isolated from cells grown in YEL at 30 °C for 16h with shaking at 200 RPM. The primer pairs used are given in Supplemental file 2. DNA: The F1/R1 primer pairs were specific for genomic sequences flanking a *hid* gene. The F2/R2 primer pair was specific for the replacement marker gene, either natR or kanR. F1/R2 and F2/R1 represent the two possible combinations of flanking-genomic and marker-specific primers. The hash symbol indicates the gene-specific primer pair corresponding to the targeted *hid*$^{+}$ allele. RNA: For the RT-PCR, the primer combinations refer to internal gene-specific primers.

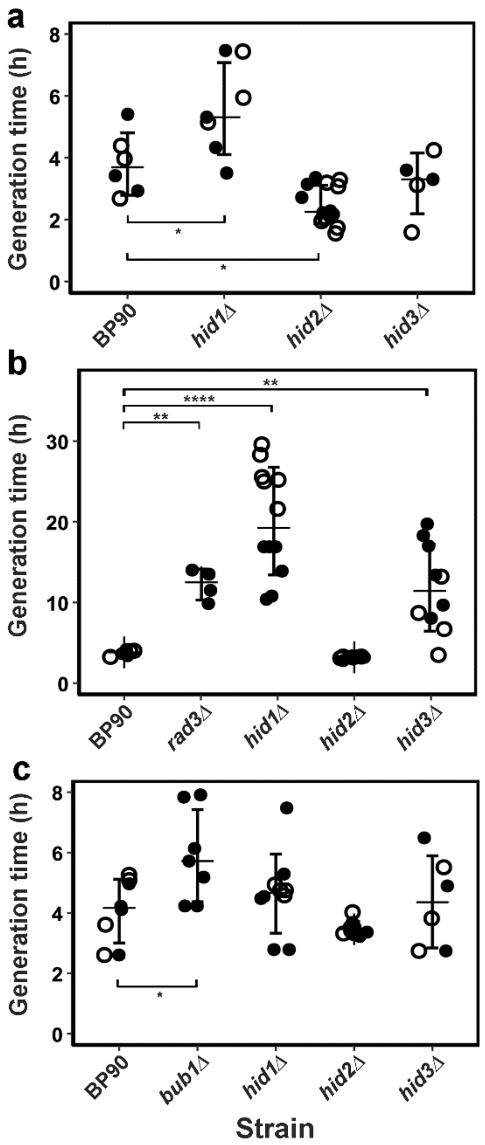


**Supplementary Fig. 5.** Effects of stress on mutants lacking Hid. Cells at an initial concentration of $5x10^5$ cell•ml$^{-1}$ were cultured in standard YEL in 48-well microtiter plates with the agent and recounted after 12 h to yield the generation time. Dark and open circles represent iso1 and iso2, respectively, for each genotype. (A) Incubation in 0.4M NaCl: *rad3Δ*, n=4; BP90, n = 8; *hid1Δ*, n= 12; *hid2Δ*, n=8; *hid3Δ*, n= 10. (B) Incubation in 0.7mM $H_2O_2$: BP90, n = 6; *hid1Δ*, n= 14; *hid2Δ*, n=10; *hid3Δ*, n= 6 (C) Incubation in 12.5 µg•ml$^{-1}$ thiabendazole (TBZ): BP90, n = 8; *bub1Δ*, n = 7; *hid1Δ*, n= 14; *hid2Δ*, n=10; *hid3Δ*, n= 6. Statistical significance: *, $p < 0.05$; ** $p < 0.01$; ****, $p < 0.0001$.

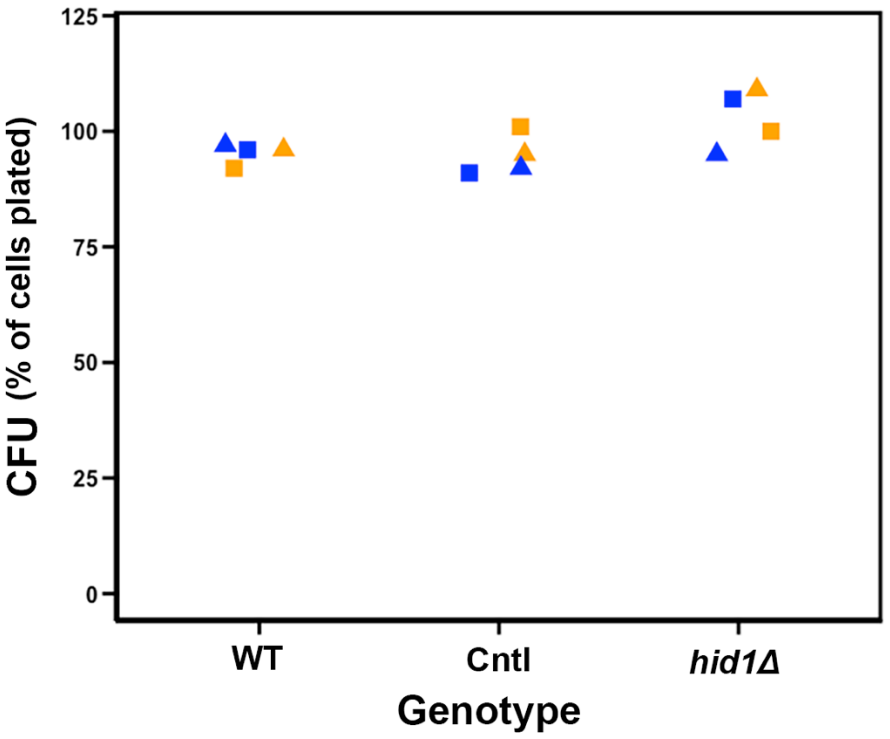


**Supplementary Fig. 6.** Estimation of relative viability of *hid1Δ* cells**.** The pre-cultures used for the proliferation analyses were counted, diluted and plated to a density of 300 colonies per petri dish on YE agar media with the appropriate auxotrophic supplements. After 60h of incubation at 30°C, the number of colonies formed were counted. WT represents the two independently propagated isolates of the multi-auxotrophic strain BP90. For each genotype, the squares and triangles represent isolates 1 and 2, respectively. The colors blue and orange represent replicates. There was no statistical difference among genotypes as indicated by single-tailed ANOVA ($F = 0.38$, $F_{crit} = 4.26$, $df = 2$).

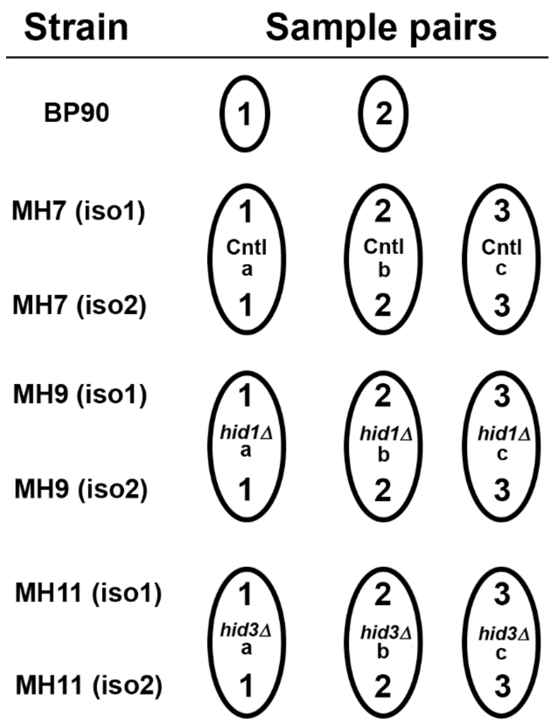


**Supplementary Fig 7.** Scheme of preparation of samples for post-genomic studies. Three independent colonies (1, 2, & 3) from each isolate were incubated in YEL plus supplements for 16h at 30$^o$ C with shaking at 200 RPM. The cells were counted and diluted to a density of 5x$10^7$ cells•$ml^{-1}$. These cultures were incubated under the same conditions until the mid-points were reached as shown in Figure 5. The cells were collected by centrifugation, washed once in sterile water, immediately frozen in liquid $N_2$, and ground to fine powder. The strains were combined at this step, where equal masses of powder were combined for the two isolates specified. The procedure yielded from 1.2 to 1.5 g of powdered cells. The circles represent the isolates and the sample numbers that were combined. As described in A third independently propagated BP90 isolate was used for the proteomic analysis.

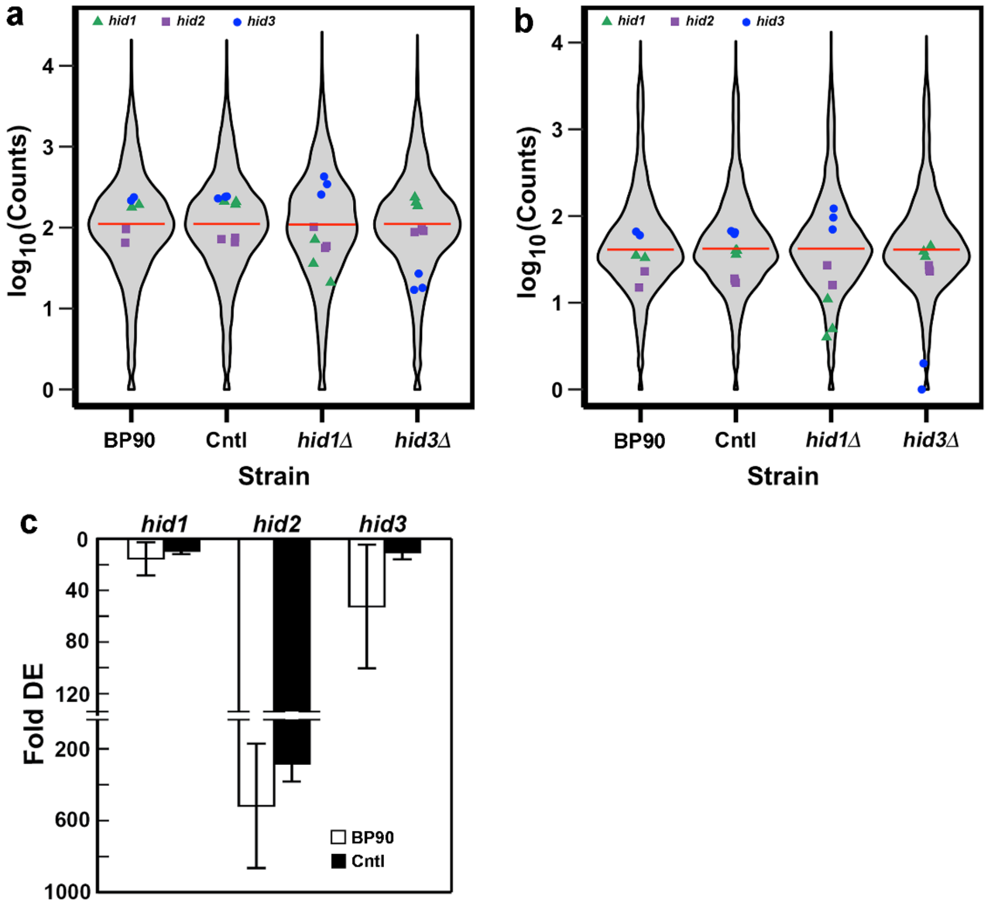


**Supplementary Fig. 8.** Relative transcript levels of *hid*+ genes**.** Transcript levels as gene Counts were extracted from the output table of the DESeq2-normalized RNA-Seq data mapped with either STAR (A) or SALMON (B) and the means for each gene calculated across samples. The points represent the counts for each *hid* gene from the individual samples. (C) Relative transcript levels determined by RT-qPCR on a Bio-rad CFX™ real-time PCR machine using Sybr green as the detection dye. For each gene in each individual sample the differential expression (Fold DE) was calculated relative to *act1*. Using the double-delta Ct method, i.e. $2^{(Ct(hid)-Ct(act1))}$. The values shown are the means ± sd (n=3) determined across the biological replicates, 2 for BP90 and 3 for Cntl. For any of the three genes, there was no significant difference between BP90 and Cntl as analysed by Student's t-test.

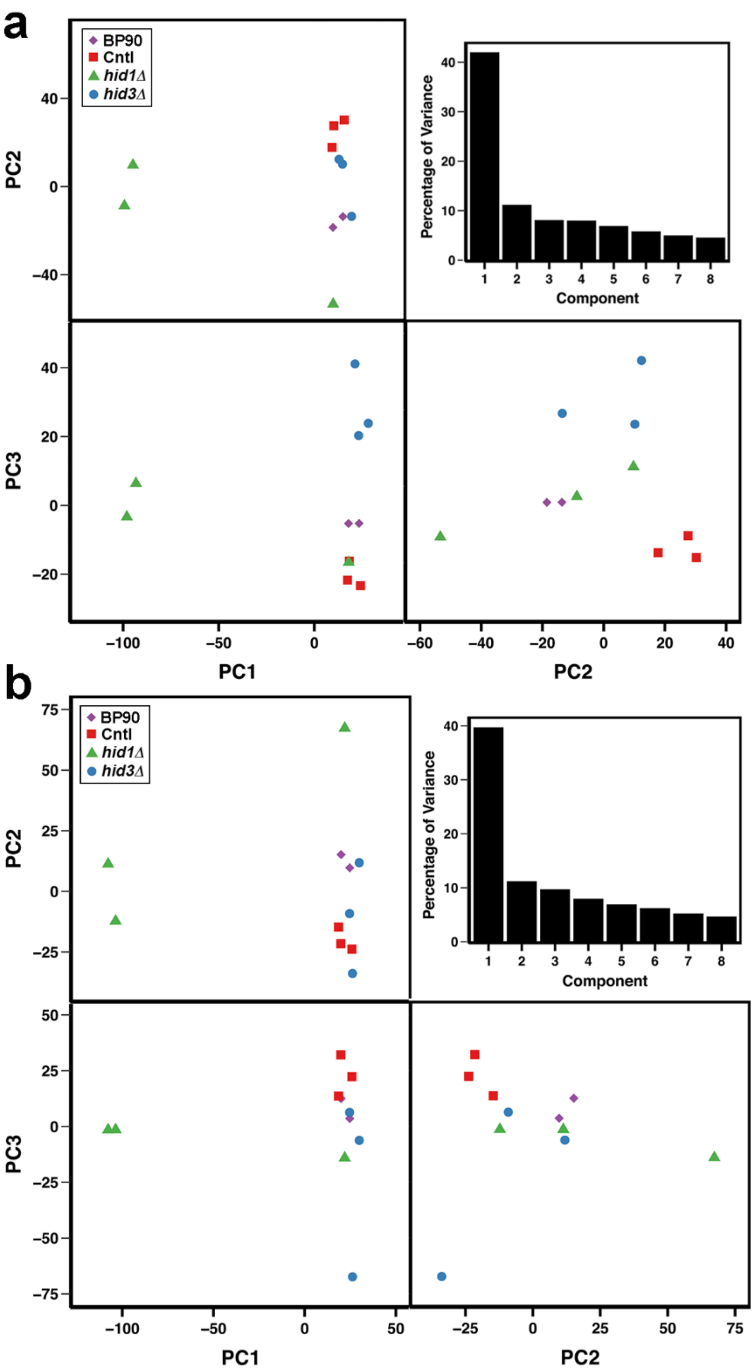


**Supplementary Fig. 9. PCA scores plot showing relationships among samples based on transcript profiling.** The counts files from STAR (a) and SALMON (b) mapping were filtered by removing all genes with a total count value of <10 across all samples. The remaining gene counts were normalized in DeSeq2 using default parameters and the PCA performed on the output file. The total variance accountable within the first 3 components was 63 % and 59 % for STAR and SALMON, respectively.

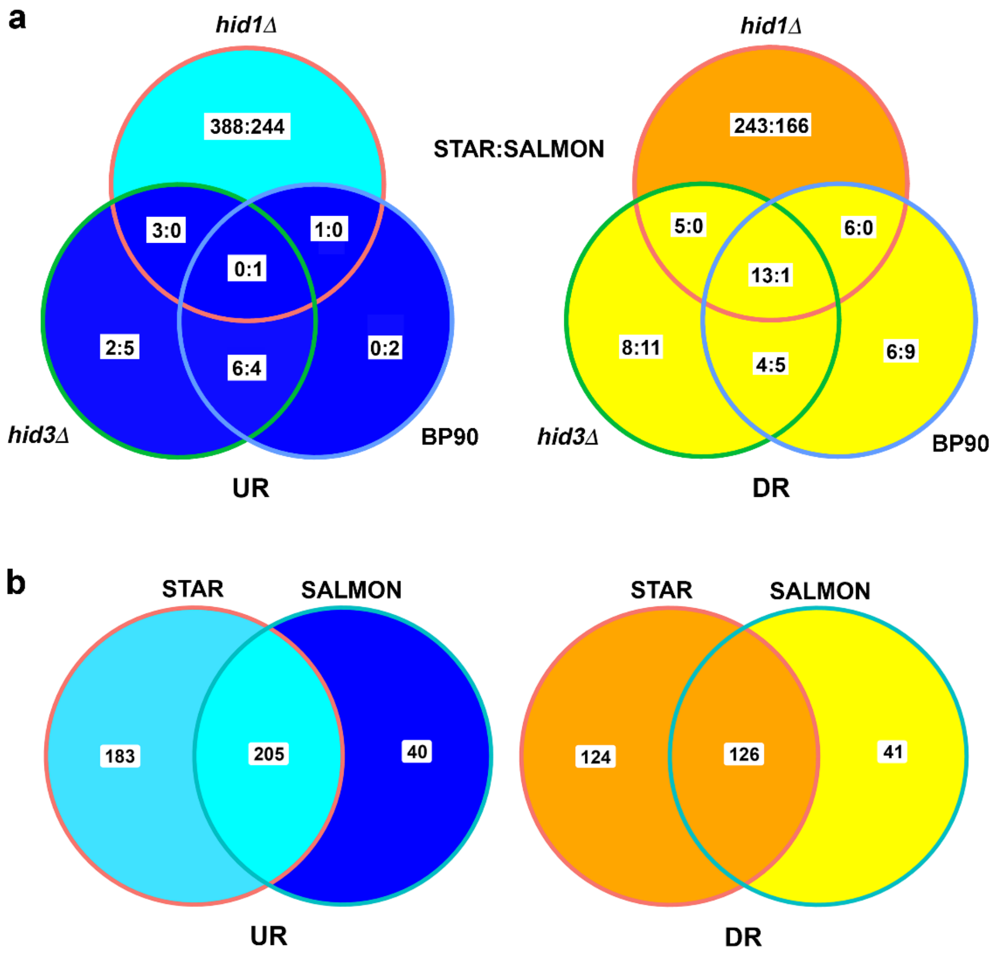


**Supplementary Fig. 10.** Venn representation of overlapping output from DESeq2 quantification of STAR and SALMON mapping. (a) The common DE genes among strains for each mapping strategy. The numbers before and after the colon represent the number of DE genes and the number in common for STAR and SALMON mapping, respectively. (b) The number of DE genes and those common between STAR and SALMON mapping strategies. The number of DE genes was determined using a p-adjusted value of 0.5 for STAR and 0.1 for SALMON.

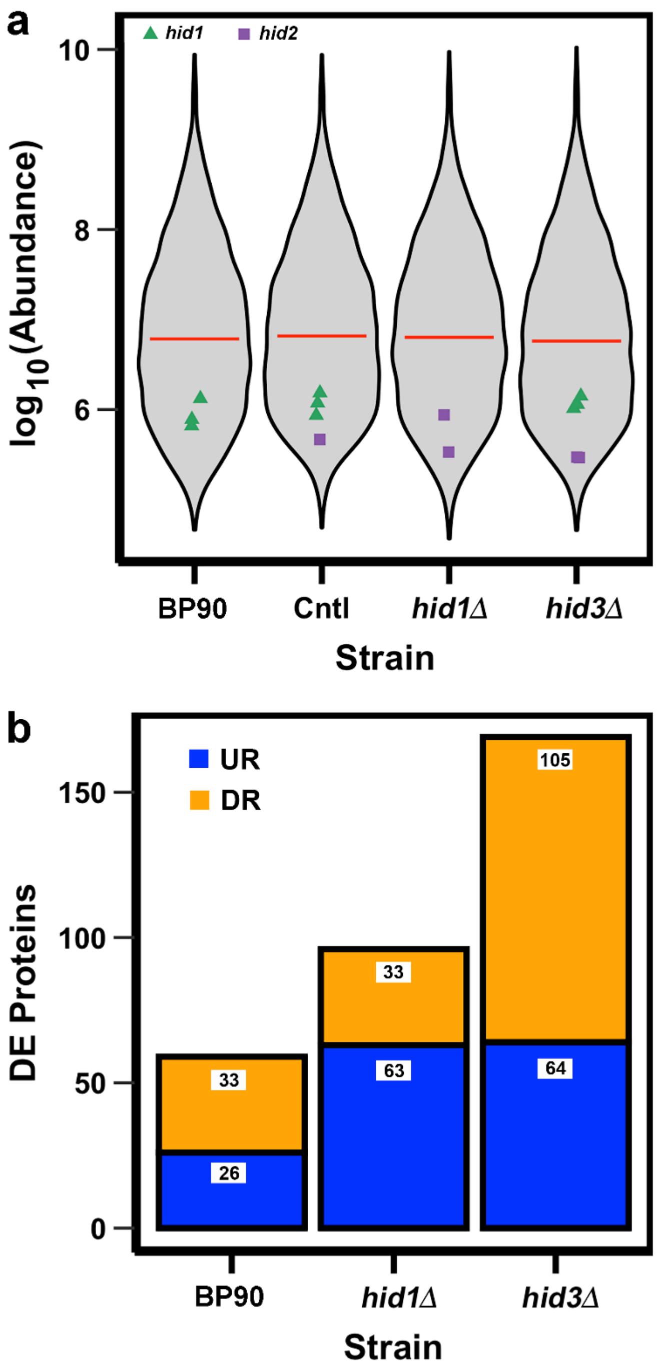


**Supplementary Fig. 11. Summary of output from proteomic analyses.** (a) Violin plots showing relative Hid protein levels. Protein levels as normalized abundances from the mass spectrometer output from the label-free quantification. The abundances of detected and quantified proteins were averaged across samples to produce the violin plots and the points represent the individual hid abundances. (b) The number of differentially expressed proteins from the label-free quantification. The numbers of proteins changing were determined by comparison to the Cntl strain. The numbers represent proteins changed in amount by more (UR) or less (DR) than an adjusted p-value $< 0.05$ as determined by ANOVA against the background median ratio and variance of all quantified proteins.

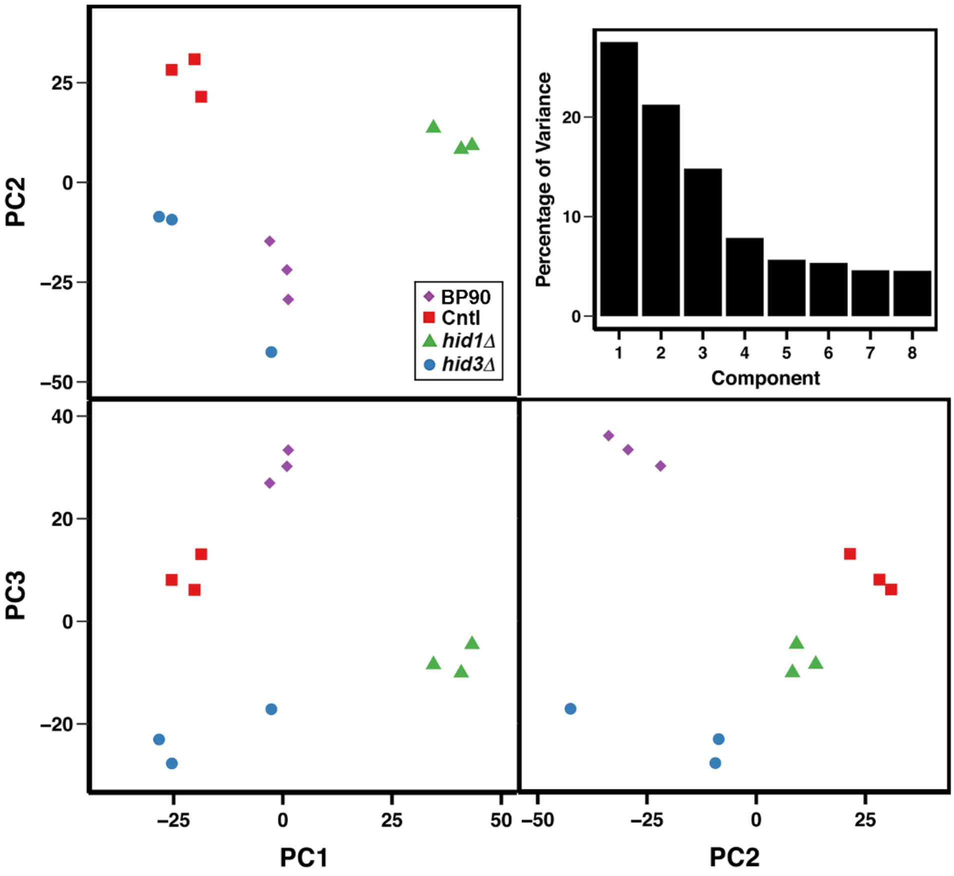


**Supplementary Fig. 12.** Principle component analysis of normalized values of quantified proteins. PCA was conducted on normalized summed peptide abundancies of unique proteins as determined by Proteome Discoverer v2.5. The number of proteins used for the PCA analysis was 2486 after removal of proteins for which more than 4 samples had values missing. Imputation of missing values for PCA was done in R using the missMDA package. PCA analysis was conducted in R using the package FactoMineR. Upper right panel shows the percentage of variance for the first eight principal components.

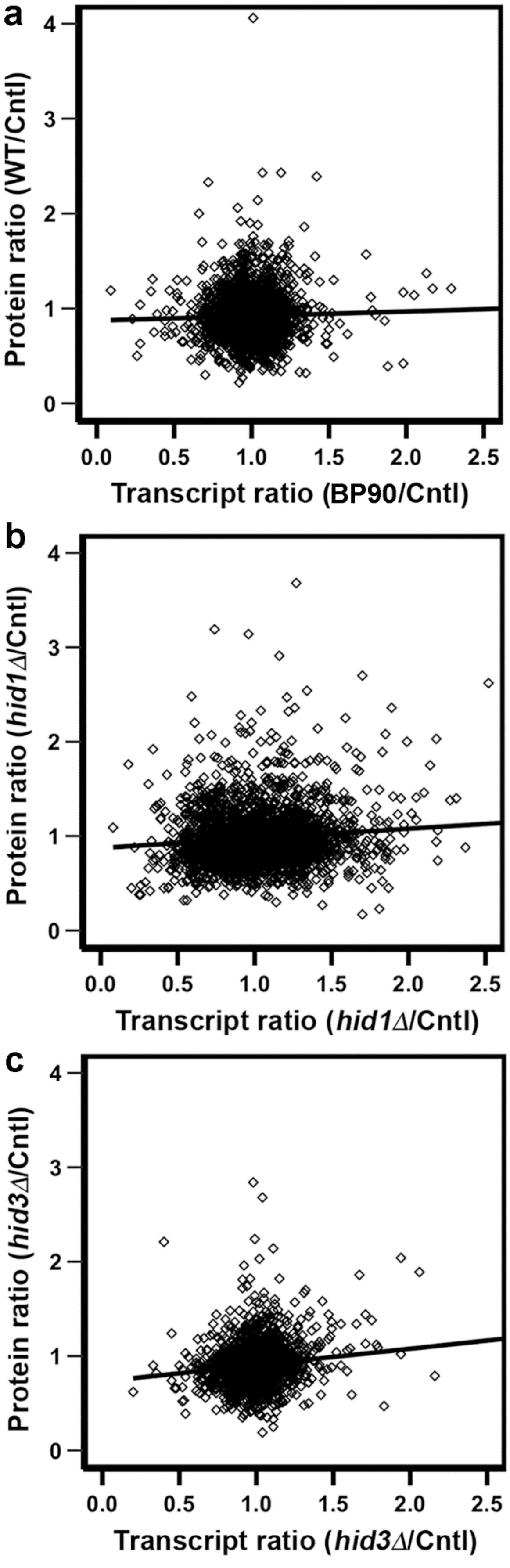


**Supplementary Fig. 13.** Correlation between gene and protein expression. The ratios used in the analysis had been pre-calculated from the analysis of differential expression of the transcriptome and proteome data as given in the additional files 5 and 6, respectively. The genes and proteins selected for the analysis were common to all three strains and represents 2244 protein entities. The line represents the best-fit Pearson linear model with the correlation coefficients given in Supplemental table 2.

**Pairwise correlation coefficients across transcriptomes and proteomes**.

| Transcript ratio | Protein ratio | r |
|---|---|---|
| BP90/Cntl | BP90/Cntl | 0.035 |
| BP90/Cntl | *hid1Δ*/Cntl | -0.054 |
| BP90/Cntl | *hid3Δ*/Cntl | 0.037 |
| *hid1Δ*/Cntl | BP90/Cntl | 0.004 |
| *hid1Δ* /Cntl | *hid1Δ*/Cntl | 0.118 |
| *hid1Δ*/Cntl | *hid3Δ*/Cntl | -0.003 |
| *hid3Δ*/Cntl | BP90/Cntl | 0.020 |
| *hid3Δ*/Cntl | *hid1Δ*/Cnt | 0.054 |
| *hid3Δ*/Cntl | *hid3Δ*/Cntl | 0.154 |
